# Signatures of relaxed selection in the *CYP8B1* gene of birds and mammals

**DOI:** 10.1101/714188

**Authors:** Sagar Sharad Shinde, Lokdeep Teekas, Sandhya Sharma, Nagarjun Vijay

## Abstract

The *CYP8B1* gene is known to catalyse reactions that determine the ratio of primary bile salts and the loss of this gene has recently been linked to lack of cholic acid in the bile of naked-mole rats, elephants and manatees using forward genomics approaches. We screened the *CYP8B1* gene sequence of more than 200 species and test for relaxation of selection along each terminal branch. The need for retaining a functional copy of the *CYP8B1* gene is established by the presence of a conserved open reading frame across most species screened in this study. Interestingly, the dietary switch from bovid to cetacean species is accompanied by an exceptional ten amino-acid extension at the C-terminal end through a single base frame-shift deletion. We also verify that the coding frame disrupting mutations previously reported in the elephant are correct, are shared by extinct *Elephantimorpha* species and coincide with the dietary switch to herbivory. Relaxation of selection in the *CYP8B1* gene of the wombat (*Vombatus ursinus*) also corresponds to drastic change in diet. In summary, our forward genomics based screen of bird and mammal species identifies recurrent changes in the selection landscape of the *CYP8B1* gene concomitant with a change in dietary lipid content.

## Introduction

The increasing number of genomes that are being sequenced across the tree of life has allowed for large-scale comparative genomic analysis. One of the most interesting findings to emerge from these analysis has been the identification of Taxonomically Restricted Genes (TRGs) and their contribution to evolutionary novelty (Albalat and Cañestro 2016; Johnson 2018). Striking phenotypic changes such as the loss of teeth in birds and the loss of vitamin C synthesis ability in primates have been convincingly linked to loss of specific genes (Hiller et al. 2012; Meredith et al. 2014). Changes in dietary patterns have also been linked to changes in digestive enzyme gene content (Wang et al. 2016; Chen and Zhao 2019; Rinker et al. 2019). These comparative approaches have immense promise in understanding the evolutionary origin of novelty. Utility of finding TRGs is best demonstrated by forward genomics approaches that have used phenotype presence/absence matrices to perform association studies that link gene loss events to specific traits in vertebrates (Sharma et al. 2018). Development of dedicated tools that improve local whole genome alignment quality have also greatly aided the search for TRGs (Sharma et al. 2016, 2017). Gene loss in multiple species with the corresponding loss of a trait in the same set of species has been used as evidence of a strong association (Hiller et al. 2012; Meredith et al. 2014). These genome-wide association studies leverage the phylogenetic relationships between species; signatures of intense relaxed selection as well as knowledge about the functional roles of these genes in coding for specific traits (see **Figure 1**). However, identification of subtle taxon-specific changes at the regulatory level continues to be challenging.

**Figure 1:**
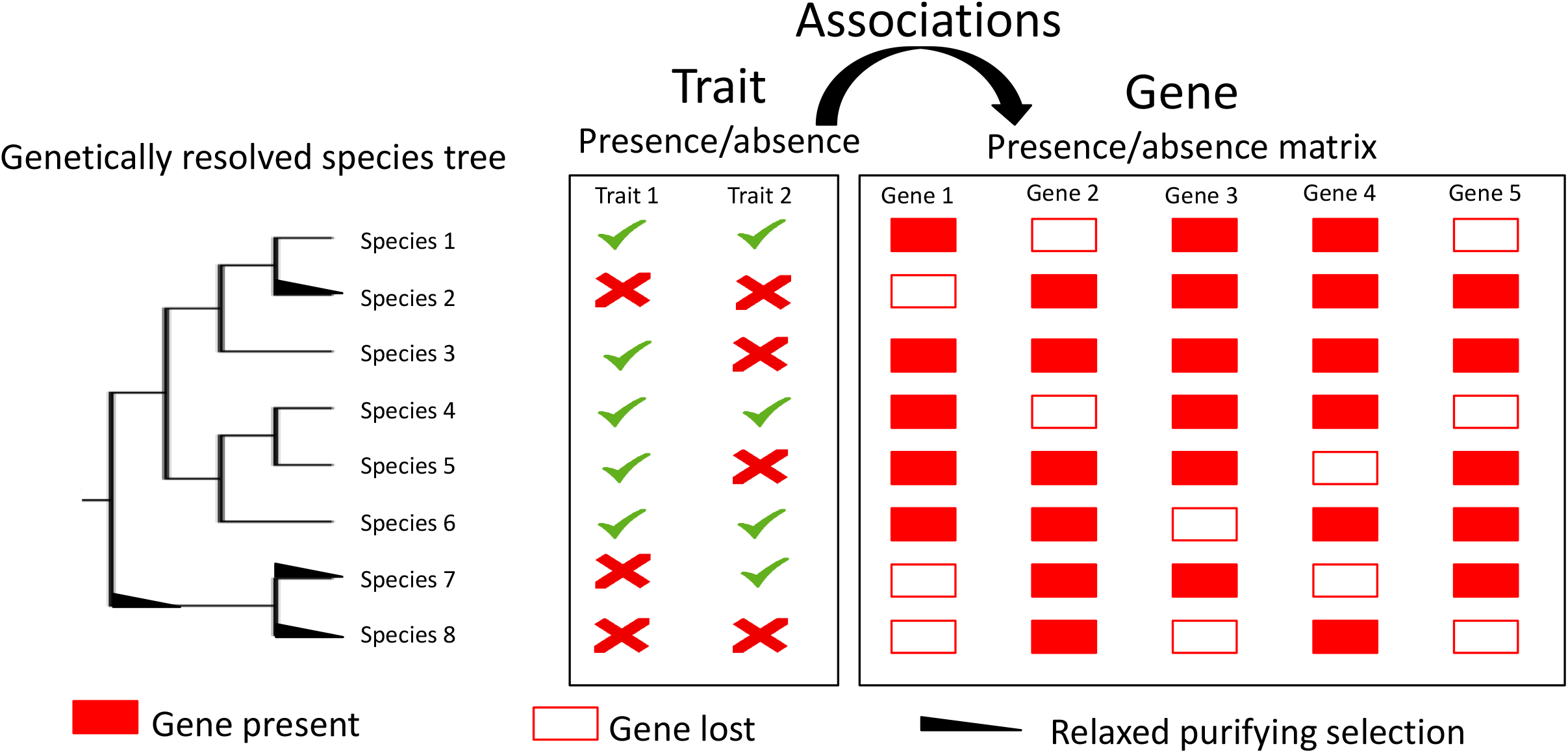
Schematic figure showing the phylogenetic relationship between species and their corresponding trait and gene presence/absence matrices. Forward genomics approaches compile such matrices using extensive bioinformatics curation and perform genome-wide associations to identify significant associations. Relaxed purifying selection at the focal gene in the foreground species (i.e., species suspected to have lost the functional copy of the gene) compared to the background species is considered as evidence for the loss of gene function. In this figure a perfect association can be seen between trait 1 and gene 1.

The bile content of animals is known to affect the digestion and absorption of lipids (Edwards 1962). Hence, genetic variation that alters the bile composition would alter lipid metabolism. The ratio of primary bile salts is determined by the reactions catalysed by the *CYP8B1* gene. Experiments in mice have shown that *CYP8B1* knockout individuals have reduced lipid absorption and increased glucose tolerance (Kaur et al. 2015; Bonde et al. 2016). Loss of this gene has recently been linked to lack of cholic acid in naked-mole rats, elephants and manatees using forward genomics approaches (Sharma and Hiller 2018). The approach pioneered by Sharma & Hiller can be used to understand the genetics of the bile pathway in diverse set of vertebrate species that have distinctive bile compositions (Hofmann et al. 2010; Hagey et al. 2010a). Identification of relaxed selection in the *CYP8B1* gene has also helped identify candidate species (example: cape golden mole) predicted to have distinctive bile composition due to the loss of genes in the bile pathway (Sharma and Hiller 2018).

In this study, we focussed entirely on the *CYP8B1* (cytochrome P450 family 8 subfamily B member 1) gene, a single exonic gene of approximately 500 amino acids that catalyses the conversion of 7 alpha-hydroxy-4-cholesten-3-one into 7-alpha, 12-alpha-dihydroxy-4-cholesten-3-one. We screened the genomes of over 200 species to look for signatures of relaxed/intensified selection in the *CYP8B1* gene that might provide clues about evolution of the bile pathway. Our screen found that (1) The *CYP8B1* open reading frame is of similar length and conserved across most bird and mammal species. Surprisingly, cetaceans have acquired a ten amino acid longer c-terminal tail through a single base pair frame-shift deletion. (2) Coding frame disrupting mutations previously reported in the *CYP8B1* gene of the elephant are correct and are shared by woolly mammoths and the American mastodon, suggesting the loss of this gene in all members of the order *Proboscidea*. (3) The previously reported signature of relaxed of selection in elephants is consistently detected irrespective of the background species used. However, we show that the detection of relaxed selection in other species of *Afrotheria* is in some cases masked by the set of background species used as well as the tree topology used. Notably, the complete open reading frame of the *CYP8B1* gene is present in the genome of the rock hyrax (a species previously reported to lack cholic acid in its bile) with no obvious signatures of relaxed selection even when different sets of background species are used. (4) A premature stop codon (AAA to TAA substitution) polymorphism is segregating in the *CYP8B1* gene of domesticated chicken. (5) We identify strong relaxation of selection in the *CYP8B1* gene of 14 other species (only 1 of these species (the wombat) shows statistically significant relaxation after correcting for multiple testing) in which the bile pathway may be disrupted or differently regulated.

## Materials and Methods

### Data and code availability

The code, data, schematic workflow and detailed instructions required to replicate the results from this manuscript are available for download under GNU license from the github repository https://github.com/ceglab/CYP8B1.

### Multiple sequence alignment and gene tree inference

The curated coding sequences from the re-annotation step (see **Supplementary Materials** for details) were used to generate a multiple sequence alignment for the *CYP8B1* gene starting from the start codon and extending until the stop codon. For species with premature stop codon, gaps in the form of N’s were inserted in place of the stop codon/frame shift inducing alleles to facilitate the alignment process. The multi-fasta open reading frames file consisting of more than 200 species was aligned through Guidance2 with codon option and 100 bootstraps independently using PRANK (v. 140603), MUSCLE (v3.8.31), MAFFT (v7.407) and CLUSTALW (2.0.12) as the alignment tool (Sela et al. 2015). To assess the consistency of the alignment across these four multiple sequence alignment tools, we compared the alignments using the SuiteMSA tool (Anderson et al. 2011). The visual comparison of the four alignments is provided as a pixel plot in **Supplementary Figure S1**. To obtain a quantitative comparison of the different alignments we also calculated the consistency between the alignments (see **Supplementary Table S1**). We found that the alignments differed from each other mainly at the alignment ends. These differences were mainly driven by the differences in the C-terminal end of cetacean species. The N-terminal end was different in two of the bat species (i.e., *Hipposideros armiger* and *Rousettus aegyptiacus*), the speckled mousebird (*Collius striatus*) and the alpaca (*Vicugna pacos*). The overall robustness of the alignment quality was deemed to be good based on the Guidance scores and presence of very few gaps. The comparison of multiple sequence alignments for each group of species was done using the program MUMSA (Lassmann and Sonnhammer 2005) to calculate the AOS (Average Overlap Score: a measure of similarity between alignments) and MOS (Multiple Overlap Score: a measure of biological correctness of individual alignments). Based on the values of AOS and MOS (provided in **Supplementary Table S2**) for each of these cases it is clear that alignment results are largely consistent across programs. We have performed tests of sequence substitution saturation described in (Xia et al. 2003) and implemented in the program DAMBE (Xia 2013). The index to measure substitution saturation (Iss) and the corresponding critical Iss (Iss.c) estimates for symmetrical and extreme asymmetrical tree are reported for each group of species that has been used in our study (**Supplementary Table S3**). In all cases we find that Iss < Iss.c and largely shows a significant difference. Hence, the sequences used in our study show little saturation.

The best sequence evolution model was identified using the program modeltest-ng (Darriba et al. 2019) with the multiple sequence alignments generated by each of the four programs as input. The best model according to BIC is provided in Supplementary Table S2. We obtained maximum likelihood gene trees for the *CYP8B1* gene using RAxML-NG with 1000 bootstraps (Kozlov et al. 2019). Subsets of these species were grouped based on their evolutionary proximity and further analyses were performed within these groups. Robustness of the results obtained was assessed using multiple tree topologies. We also evaluated the sequences for substitution saturation using the index to measure substitution saturation (Iss) described in (Xia et al. 2003) and implemented in the program DAMBE (Xia 2013). The index to measure substitution saturation (Iss) and the corresponding critical Iss (Iss.c) estimates for symmetrical and extreme asymmetrical tree are reported for each group of species that has been used in our study (**Supplementary Table S3**). In all cases we find that Iss < Iss.c and largely shows a significant difference. Hence, the sequences used in our study show little saturation.

The nucleotide composition bias across the *CYP8B1* gene sequence was calculated for each species. The mean GC content is reported on the github page: (https://github.com/ceglab/CYP8B1/ORFs/README.md. The mean GC content values range from 48% to 66%, with the highest values of GC content being found within Chiroptera and the Platypus. We have also investigated the heterogeneity in GC content along the length of each sequence by calculating the GC content in sliding windows of 100 base pairs with a step size of 10 base pairs (https://github.com/ceglab/CYP8B1/gc_content). The heterogeneity in GC deviation and GC content across the gene sequence for each of the species can be seen in **Supplementary Figure S2A** and **S2B** respectively.

### Test for relaxed or intensified selection and episodic diversifying selection

The test for relaxed or intensified selection implemented in the RELAX program compares a background set of species with a foreground set of species to detect signatures of relaxation in selection or intensification of selection in a hypothesis testing framework (Wertheim et al. 2015). Here, we have used *CYP8B1* gene sequences from more than 200 species to look for signatures of relaxed selection in subgroups using the program RELAX available in the package HyPhy (hypothesis testing using phylogenies) (Pond et al. 2005). While a minimum of 1 foreground species can be tested using RELAX, at least two other species need to be included in the alignment with at least one of them being labelled as a background species. The tests for selection were performed using the aBSREL (Kosakovsky Pond et al. 2011; Smith et al. 2015), MEME (Murrell et al. 2012), FEL (Kosakovsky Pond and Frost 2005), FUBAR (Murrell et al. 2013) and BUSTED (Murrell et al. 2015) methods implemented in the HyPhy package (Smith et al. 2015).

## Results

Intense signatures of relaxed selection have generally been found in genes that have acquired reading frame disrupting changes and are considered as further evidence supporting gene loss (Sharma and Hiller 2018). Nonetheless, relaxed selection can also have other consequences such as phenotypic plasticity and need not always lead to gene loss (Lahti et al. 2009). We have performed manual curation of the *CYP8B1* gene to correct genome assembly and annotation artefacts in numerous species (see **Supplementary Materials**). These curated open reading frames are used to test for signatures of relaxed selection in more than 200 species.

### Cetacean specific c-terminal peptide tail

Our dataset consists of *CYP8B1* gene sequences from sixteen cetacean species. We found that the open reading frame of all cetacean species except those from the genus *Sousa (Sousa chinensis* and *Sousa sahulensis)* and *Tursiops (Tursiops truncates* and *Tursiops aduncus)* were approximately 30 base pairs longer, leading to an additional 10 amino-acid peptide tail that is missing in all the other artiodactyl species (see **Figure 2A**). This 10 amino-acid stretch was conserved across twelve cetacean species and was also supported by raw sequencing reads (see https://github.com/ceglab/CYP8B1/SAMs). To check whether this is a cetacean specific change, we added the *CYP8B1* gene sequence from ten non-cetacean artiodactyl species to our multi-species sequence alignment and found that all cetacean species had acquired a one base-pair deletion exactly three codons (CCT to C-T) before the erstwhile stop codon found in non-cetacean artiodactyl species (see **Figure 2B**). This single base-pair deletion resulted in a coding frame shift and a downstream stop codon after the split of the cetacean ancestor from the hippopotamus.

**Figure 2:**
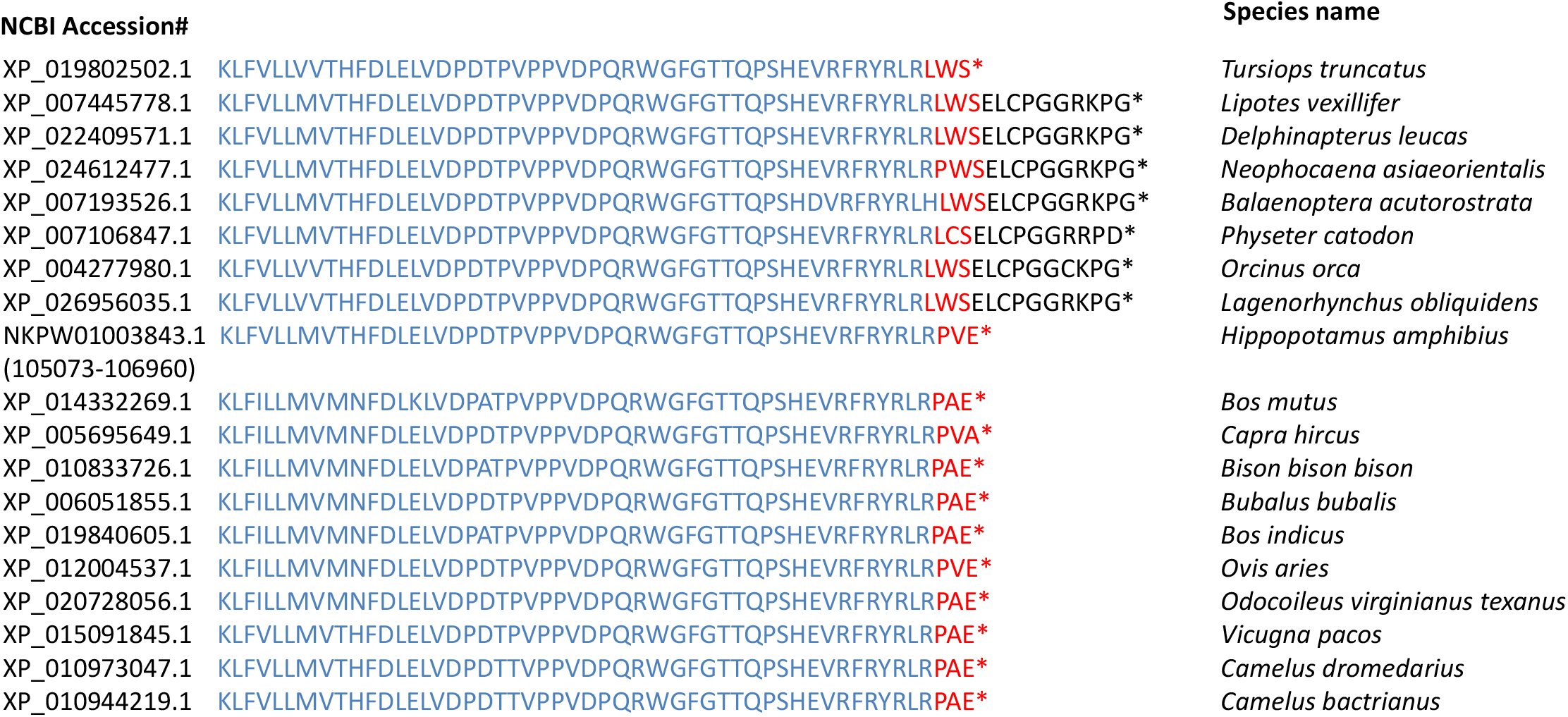
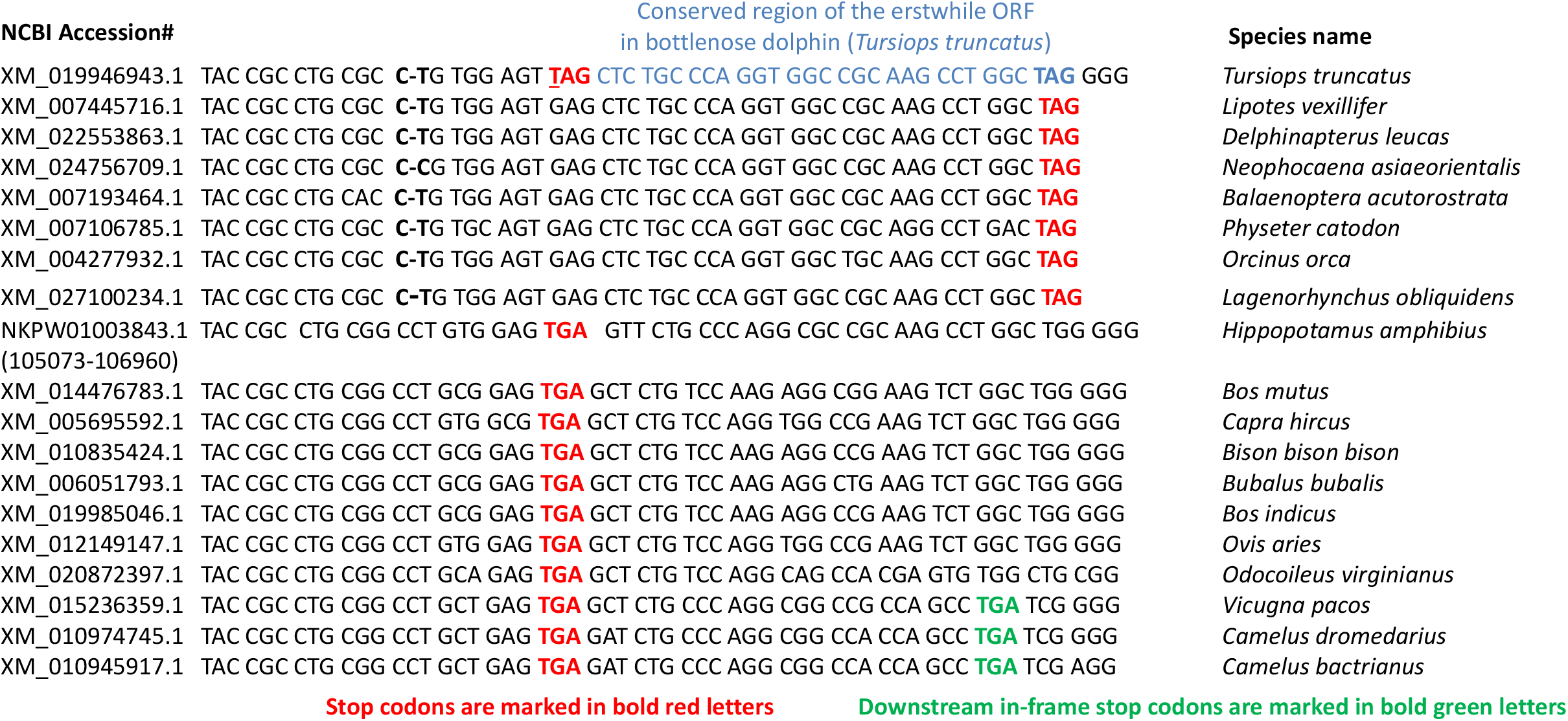
Multiple sequence alignment of cetacean and non-cetacean Artiodactyl gene sequences for the *CYP8B1* gene. (A) Ten amino-acid extension at C-terminal of cetaceans (B) Nucleotide sequence alignment showing the one base-pair frame-shift deletion across all cetaceans followed by a single stop-codon inducing transversion in the bottlenose dolphin (*Tursiops truncatus*).

Lipid metabolism genes are known to have evolved under positive selection in the cetacean lineage (Foote et al. 2015, 2016; Wang et al. 2016). A more detailed study of the lipid metabolism genes in a recent study found that the *CYP8B1* gene has the strongest pattern of positive selection in cetaceans (Endo et al. 2018). The single base pair deletion in the *CYP8B1* of cetaceans was not identified in previous studies to the best of our knowledge. However, the co-occurrence of gene-damaging indels, positive selection and dietary changes has been reported for the *CYP3A* gene cluster in humans (Hu and Ng 2012; Wagh et al. 2012). Gene-damaging indels are known to be accompanied by compensatory changes that maintain the complete reading frame. Frame shift inducing in-del changes can drastically alter the protein sequence through even a single base insertion or deletion. The ten amino-acid change in protein length of the *CYP8B1* gene in cetaceans is intriguing given the central role of this gene in the bile pathway. The change from a herbivorous diet in the bovid ancestor to a carnivorous diet in cetaceans is also accompanied by positive selection in the *SLC27A2* and *HSD17B4* genes that are part of the bile pathway (Endo et al. 2018). We have used our alignment to identify the sites under selection in the cetacean lineage (see **Supplementary Table S5**). Hence, the *CYP8B1* gene shows signatures of episodic positive selection at few sites.

The bottlenose dolphin (*Tursiops truncatus*) stop codon has been acquired subsequently through an independent (G to T) transversion event that converts the GAG codon to a TAG stop codon (see **Figure 2B**). The recent (after the split from the pacific white sided dolphin (*Lagenorhynchus obliquidens*)) independent change in coding region length in bottlenose dolphin is further supported by the strong conservation of the region downstream from the new stop codon until the old stop codon. Dolphins have a bile profile that is more similar (consisting of CA and CDCA) to the hippopotamus and bovid species in contrast to whales that have a distinctive bile profile consisting of DCA, CA and CDCA (Hofmann et al. 2010). However, it is not clear whether this distinctive bile composition has a genetic basis. Nonetheless, the repeated change in the length of the C-terminal peptide tail of the *CYP8B1* gene needs to be investigated further to understand if it has had any functional consequences. The multi-species alignment spanning cetaceans was analysed using the general descriptive model implemented in RELAX. We did not use the information obtained from the general descriptive model for performing any other tests as that would amount to using the same data twice. Although none of the cetacean species showed (see **Supplementary Table S4**) significant intensification or relaxation, it can be seen (**Supplementary Figure S3A**) that the ancestor of all cetaceans experienced relaxed selection (K<1) coinciding with the loss of the stop codon through a one base-pair deletion. Moreover, the intermittent relaxed selection (K<1) within various cetacean lineages is in sharp contrast to the presence of intensified selection (K>1) in most non-cetacean artiodactyl species. A previous study has already used tests for positive selection comparing bovid species with cetaceans and has found evidence for positive selection in the lineage leading to cetaceans (Endo et al. 2018). Hence, it seems that the C-terminal tail of the *CYP8B1* gene underwent changes in length potentially as a result of intermittent relaxation in purifying selection accompanied by episodic positive selection.

### Gene loss and relaxed selection within *Afrotheria*

All the five mutations that cause loss of the *CYP8B1* gene are known to be shared between the African and Asian elephant, suggesting the lack of cholic acid in the common ancestor of both elephant species (Sharma and Hiller 2018). We further screened the genomic reads of the woolly mammoth (*Mammuthus primigenius*), Columbian mammoth (*Mammuthus columbi*), straight-tusked elephant (*Palaeoloxodon antiquus*) and the American mastodon (*Mammut americanum*) and found that all the five coding frame disrupting changes were shared between the woolly mammoth, American mastodon and the African elephant (see **Supplementary Table S6**). Despite the poor read coverage and potential DNA damage induced artefacts, both the Columbian mammoth (*Mammuthus columbi*) and straight-tusked elephant (*Palaeoloxodon antiquus*) showed evidence for few of the coding frame disrupting changes (see **Supplementary Table S6**).

The West Indian manatee (*Trichechus manatus*) shares only one of the five coding frame disrupting changes with the elephant lineage. Hence, the remaining four changes should have occurred after the ancestor of the lineage leading to mastodons, elephants and woolly mammoths split from the ancestor of manatees. However, the possibility of independent loss of the *CYP8B1* gene in these two groups cannot be ruled out. Based on dental wear and distribution of vegetation, proboscideans are thought to have had a herbivorous diet (Saarinen and Lister 2016). Similar to the dietary switch between bovids and cetaceans, the change in selection landscape within Afrotheria coincides with changes in diet.

We used seven Afrotherian species and performed independent tests for relaxed selection in each of the species (treating one species as the foreground each time) and the remaining species as the background. A significant relaxation of selection (K<1 at p-value <0.01) was found in *Loxodonta africana* and strong intensification of selection in the lesser hedgehog tenrec. While significant relaxed selection has also been reported for *Trichechus manatus latirostris* and *Chrysochloris asiatica* in a previous study (Sharma and Hiller 2018), we found that neither of these two species showed significant relaxed selection when the six species used in our alignment were used as the background species (see **Supplementary Table S4**). Since the set of species used as the background was different in our test compared to the previous report we systematically investigated the effect of using different species as the background as well as different tree topologies.

### Effect of background species choice and tree topology on tests of relaxed selection

In order to evaluate the effect of using different background species in our tests of relaxed selection we used all combinations of background species that are possible within our seven Afrotherian species and performed independent tests of relaxed selection for each of the species. We have performed tests of relaxed selection using all possible combinations of background and foreground species using not just the strict consensus tree but using each of the topologies (topology 1 to 3, see **Figure 3A**) that were supported by at least 10 trees out of the 1000 bootstraps performed. As a contrast to comparing the 3 most well supported topologies, we also evaluated 3 randomly chosen trees that are supported by only 1 tree out of 1000 (topology 4 to 6, see **Figure 3A**). Species that are neither labelled as foreground nor background are treated as unclassified. To evaluate the effect of only changing the species labelling, the same alignment was used for all of these tests. Consistent with previous reports, we found that tests for relaxed selection in *Loxodonta africana, Trichechus manatus latirostris* and *Chrysochloris asiatica* were significant (without correcting for multiple testing) when the background set did not contain species under relaxed selection (see **Figure 3B**). Among the species showing signatures of relaxed selection, the results of tests in the cape golden mole (*Chrysochloris asiatica*) seem to be affected the most by the tree topology.

**Figure 3:**
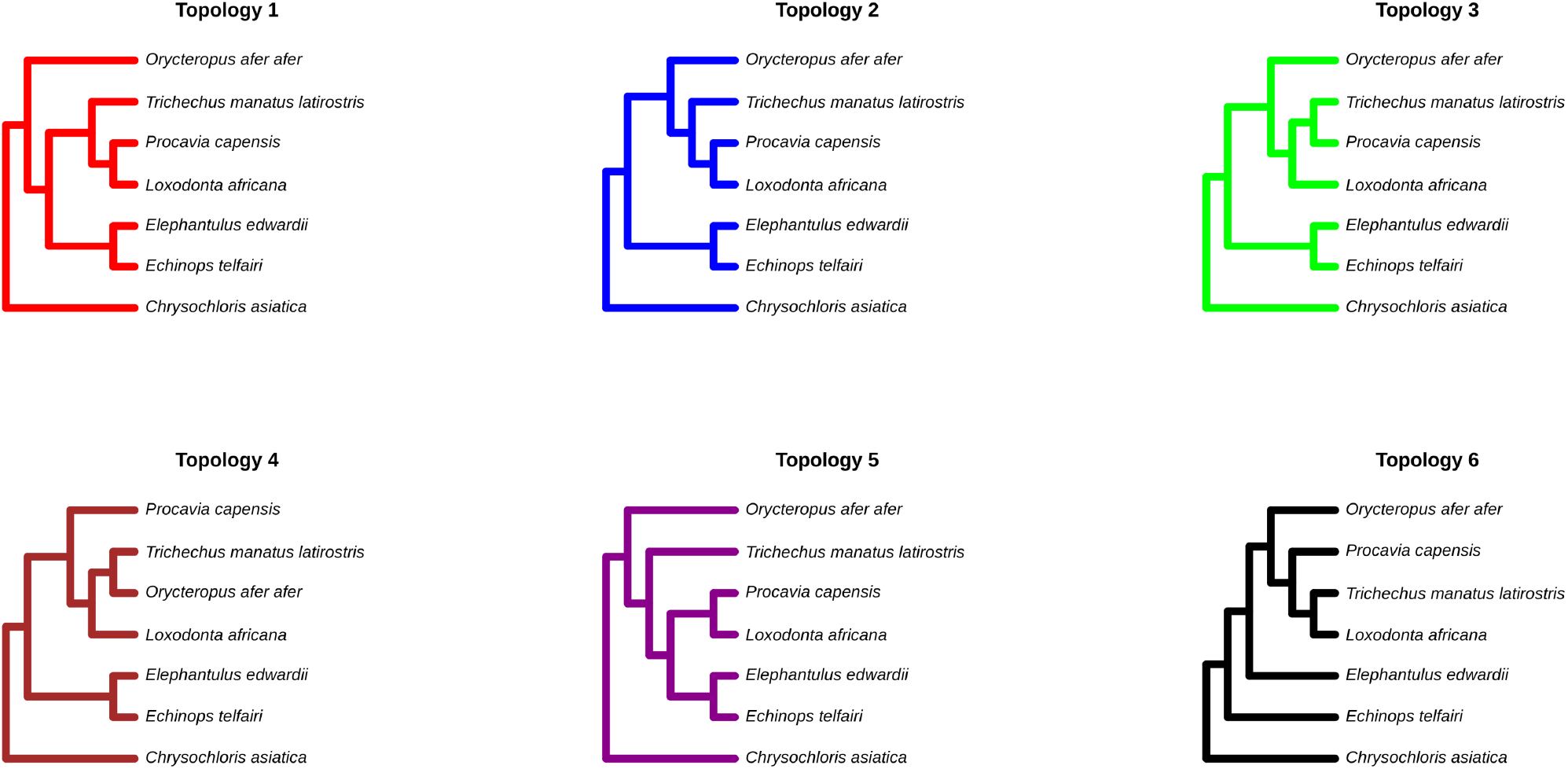
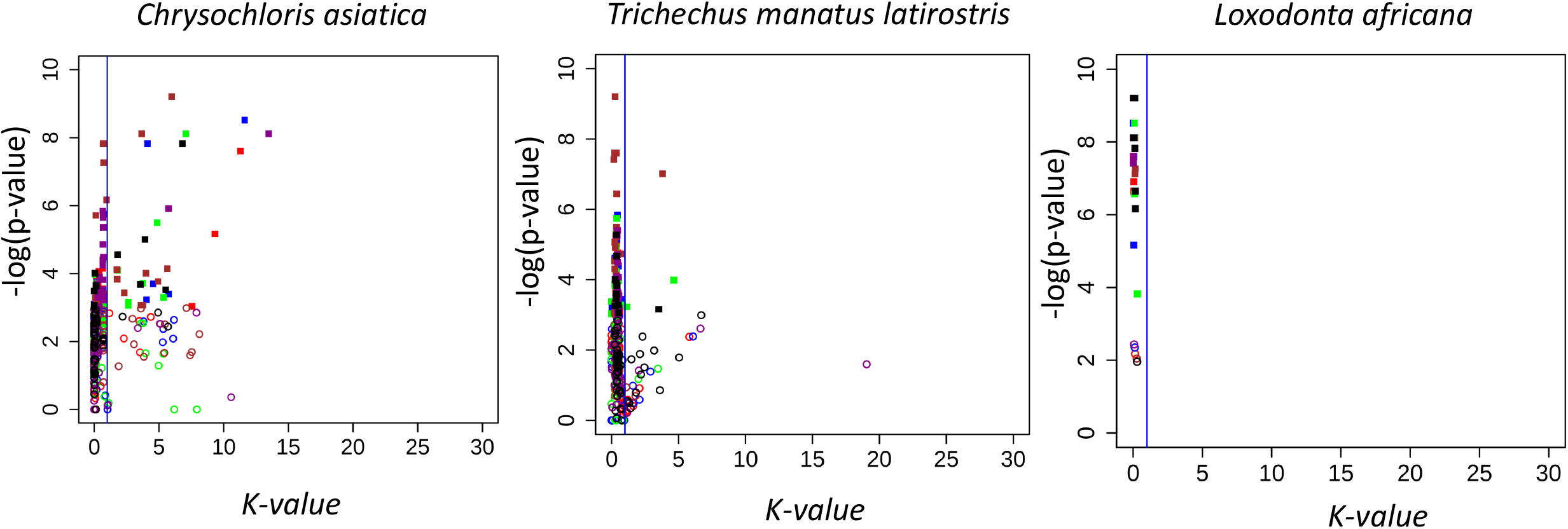
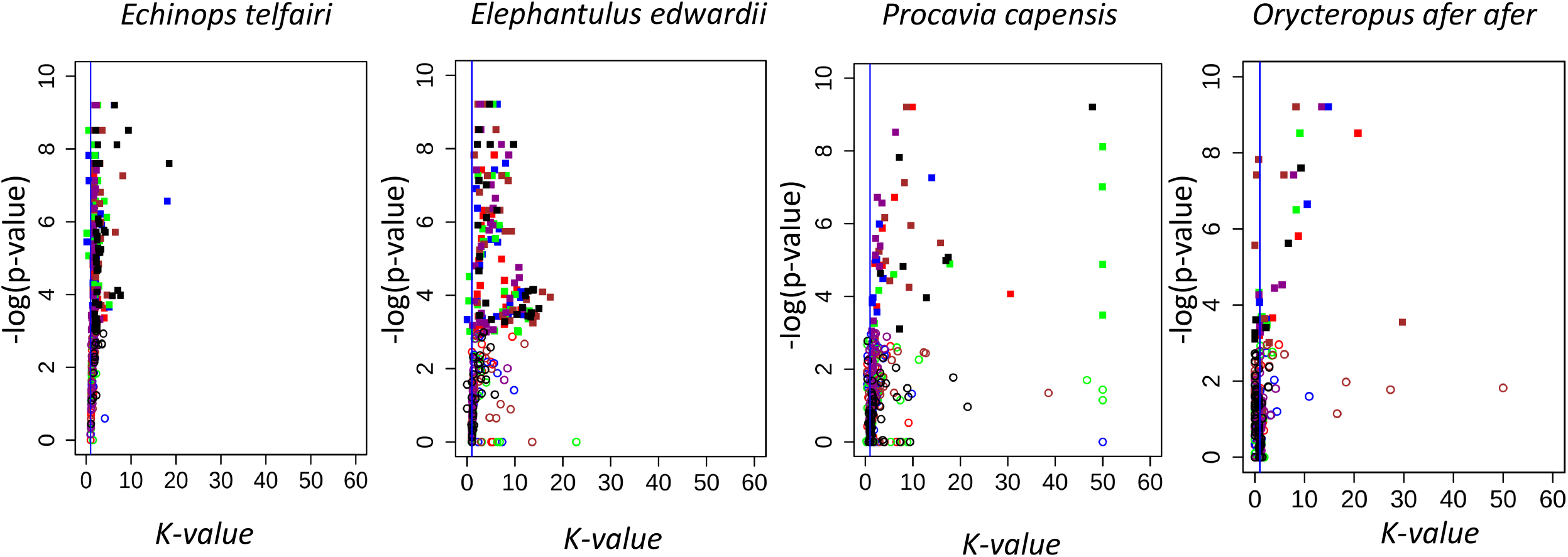
Signatures of relaxed selection represented as a combination of K values (K<1 is relaxed selection & K>1 intensified selection) denoting the intensity of change in selection and corresponding p-values in species from the order *Afrotheria*. Each data point in the figure is the result of one hypothesis test each. Tests that showed significant relaxation of selection are shown in filled squares while those that are not significant are shown as empty circles. The colour of each data point corresponds to one of the six tree topologies shown in panel A. (A) Tree topologies used to assess signatures of relaxed selection (B) Species that consistently showed significant relaxed selection (C) Species that consistently showed significant intensified selection.

Despite using a varied set of background species we found that the rock hyrax (*Procavia capensis*), lesser hedgehog tenrec (*Echinops telfairi*), cape elephant shrew (*Elephantulus edwardii*) and the aardvark (*Orycteropus afer afer*) showed signatures of intensified selection (see **Figure 3C**). Consistent with the results obtained from the above analysis, we found that the general descriptive model also shows strong relaxation in the elephant and manatee lineages (**Supplementary Figure S3B**) and intensification in the lesser hedgehog tenrec. Based on these extensive empirical analyses with different tree topologies, we believe that while tree topology does have an effect on the results, it is much less of a concern than the set of species used as the background and foreground set.

We performed tests for relaxation/intensification of selection in each of the ten species from the order *Chiroptera* considering the remaining nine species as the background set. Strong intensification of selection (**Supplementary Table S7**) was seen in the Natal long-fingered bat (*Miniopterus natalensis*) and the black flying fox (*Pteropus alecto*). However, tests in three out of the ten species resulted in highly unreliable estimates of K. To ensure that these tests are not strongly affected by the set of species used as the background, we performed additional tests for relaxation/intensification of selection in each of the species after removing one of the ten species from the alignment. We found that the large flying fox (*Pteropus vampyrus*) and the great roundleaf bat (*Hipposideros armiger*) showed relaxed selection in most of the tests when one of background species was removed (**Supplementary Table S7**). The Natal long-fingered bat (*Miniopterus natalensis*) continued to show intensification of selection even when one of the bat species was removed from the alignment. These patterns seen in Afrotheria and *Chiroptera* highlight the relevance of background species used for hypothesis testing.

### Signatures of relaxed selection in Rodents, Bears, Lemurs and the Wombat

Loss of the *CYP8B1* gene in naked-mole rat (*Heterocephalus glaber*) has been shown by the conserved synteny of the flanking regions, changes in the length of intergenic regions and lack of supporting reads in sequencing datasets (Sharma and Hiller 2018). Among the rodent species with genome assemblies, the Damara mole rat (*Fukomys damarensis*) is evolutionarily the closest to the naked mole rat. Our study finds signatures of relaxed selection (see **Supplementary Table S4**) in the Damara mole rat and provides additional support for the inferred gene loss previously reported in the naked mole rat. Both the naked-mole rat (*Heterocephalus glaber*) and the Damara mole rat (*Fukomys damarensis*) are herbivorous and represent a shift in diet from other rodents. Based on the species included in our alignment, we could see relaxed selection in *Mus pahari* and *Mus spretus* and intensified selection in *Mus musculus* and *Mus caroli* (**Supplementary Table S4**). Previous studies have reported the presence of only β-MCA, α-MCA and CDCA in the African pygmy mouse (*Mus minutoides*) and only β-MCA, CA and α-MCA in the South eastern Asian house mouse (Mus *musculus castaneus*). Such rapid change in the bile composition and patterns of relaxed/intensified selection suggests divergence of bile pathway might have played an important role in the diversification of the *Mus* subgenus and potentially reflects rapid changes in dietary patterns.

The diet of the North American beaver (*Castor canadensis*) mainly consists of tree bark and cambium (the soft tissue that grows under the bark) and its bile composition has been shown to contain only UDCA and CDCA (Hagey et al. 2010b). It is possible to enzymatically obtain UDCA starting from CDCA or CA or even lithocholic acid for industrial production (Eggert et al. 2014; Tonin and Arends 2018). Hence, it is intriguing to evaluate whether the *CYP8B1* gene is still required in the beaver genome as it is possible produce both CDCA and UDCA without having to produce cholic acid. Based on manual curation and correction of the beaver *CYP8B1* gene sequence we could annotate a complete open reading frame that is also supported by raw reads (see https://github.com/ceglab/CYP8B1/SAMs). However, we could find relaxation of selection in the beaver lineage (see **Supplementary Table S4**). The relative contribution of CA vs CDCA to the formation of UDCA can help understand the need for retaining a functional copy of the *CYP8B1* gene in the beaver genome and the patterns of relaxed selection identified.

It has previously been reported that the bile composition of the family *Lemuridae* consists mostly of CDCA (Hagey et al. 2010b). We used an alignment of six species consisting of *Prosimians* & *Dermoptera* to look for signatures of relaxed selection. Strong relaxed selection (see **Supplementary Table S4**) was inferred in the black lemur (*Eulemur macaco*) and the Philippine tarsier (*Carlito syrichta*). This raises the intriguing possibility of complete lack of cholic acid in lemurs through disruption of the bile pathway. However, the Philippine tarsier is known to have cholic acid in its bile (Hagey et al. 2010b). While signatures of relaxed selection can help identify candidate species, high quality genomes are required to identify gene disrupting mutations that might be present in one of the bile pathway genes.

Based on an alignment of six species spanning *Monotremata, Marsupialia* and *Xenarthra* we found evidence for significant (even after correcting for multiple testing) relaxed selection (see **Supplementary Table S4**) in the wombat (*Vombatus ursinus*). In contrast to this, significant intensified selection was seen in the Queensland koala (*Phascolarctos cinereus*). Multiple copies of varying lengths have been assembled and annotated for the *CYP8B1* gene in the Tasmanian devil (*Sarcophilus harrisii*) and opossum (*Monodelphis domestica*). Hence, we did not include these two species in our alignment. It has been reported that the wombat has CDCA and unusual 15α-OH bile acids as the major bile components (Hagey et al. 2010b) and might explain the strong relaxation of selection in its *CYP8B1* gene. Moreover, wombats are herbivorous and represent a dietary shift. We could not see any evidence for the presence of multiple copies of the *CYP8B1* gene within the wombat in the raw sequencing read data (see https://github.com/ceglab/CYP8B1/SAMs). The significant intensified selection seen in the koala is potentially reflective of the Queensland koala (*Phascolarctos cinereus*) having a bile composition consisting almost entirely of oxoLCA (Hagey et al. 2010b) and a diet that mainly consists of eucalyptus leaves. The gene-wide test for branch specific selection implemented in the program BUSTED (Murrell et al. 2015) also showed evidence of significant (p-value < 0.01) episodic diversifying selection in the koala branch. However, the signatures of positive selection seen in the koala lineage were not significant after removal of the wombat sequence from the alignment.

### Selection landscape in bird lineages

The bile composition across birds has considerable diversity (Hofmann et al. 2010; Hagey et al. 2010a). Hence, we hoped to find distinctive molecular signatures reflecting the previously reported bile composition. Our expectation is also supported by recently reported patterns of molecular evolution in other dietary enzymes in birds (Chen and Zhao 2019). We analysed different clades of birds by grouping them as *Galliformes*, Ducks, *Palaeognathae, Passeriformes, Telluraves* (without passerines) and *Aequorlitornithes* (see Supplementary Material for details).

### Population gene dispensability in chicken

The chicken genome assembly is a good starting point within birds for performing gene gain/loss analysis as it has the oldest and potentially the most well curated and continually updated genome assembly among birds (International Chicken Genome Sequencing Consortium 2004). Moreover, the latest build of the chicken genome Gallus_gallus-5.0 has been assembled to a high quality using long read sequencing methods (Warren et al. 2017). To maintain consistency with the previous assemblies, the same highly inbred red jungle fowl individual used for the initial genome sequencing effort has been used in the recent chicken genome assembly (Warren et al. 2017). The reference genome individual and its associated raw read data provide compelling evidence for a premature stop codon (AAA to TAA) in the UCD-001 individual that results in a 13% reduction in the amino-acid sequence length of the *CYP8B1* gene (see **Supplementary Table S10**).

Chicken bile salts are known to contain cholic acid and the inferred premature stop codon/gene loss event appears contradictory. To verify whether this substitution is segregating as a polymorphism in chicken breeds, we screened the re-sequenced whole genomes of more than 100 chicken individuals to look for this stop codon and find that it is found as heterozygous (TAA) in at least three individuals and homozygous (TAA/AAA) in another individual (see **Figure 4** and **Supplementary Table S10**). The *CYP8B1* gene open reading frame was found to be conserved across bird species and none of the other bird species had a stop codon at the position that the chicken *CYP8B1* gene has acquired a stop codon. We found the ancestral codon state (AAA) in the one *Gallus varius* individual screened. This suggests that this premature stop codon inducing polymorphism was potentially acquired in domesticated chicken. To further verify whether the *CYP8B1* gene is lost in chicken due to relaxed selective constraints, we decided to use the strategy used by (Sharma and Hiller 2018) and looked for signatures of relaxed selection in the chicken lineage. We found evidence for relaxation of selection in the chicken lineage when the *CYP8B1* sequence of the greater prairie chicken (*Tympanuchus cupido pinnatus*) was included in the alignment (see **Supplementary Table S4**). To further evaluate the robustness of this result, we used three of the most well supported tree topologies (see **Supplementary Figure S4A**) to test for signatures of relaxed selection while using all possible combination of species as the background set for each of the six species included (see **Supplementary Figure S4B, S4C**). We found consistent signatures of relaxed selection in chicken (see **Supplementary Figure S3C**). Loss of the *CYP8B1* gene has been shown to have beneficial consequences (Kaur et al. 2015) and a premature stop codon could have been selected during the domestication process in some breeds while being under relaxed purifying selection in other breeds.

**Figure 4:**
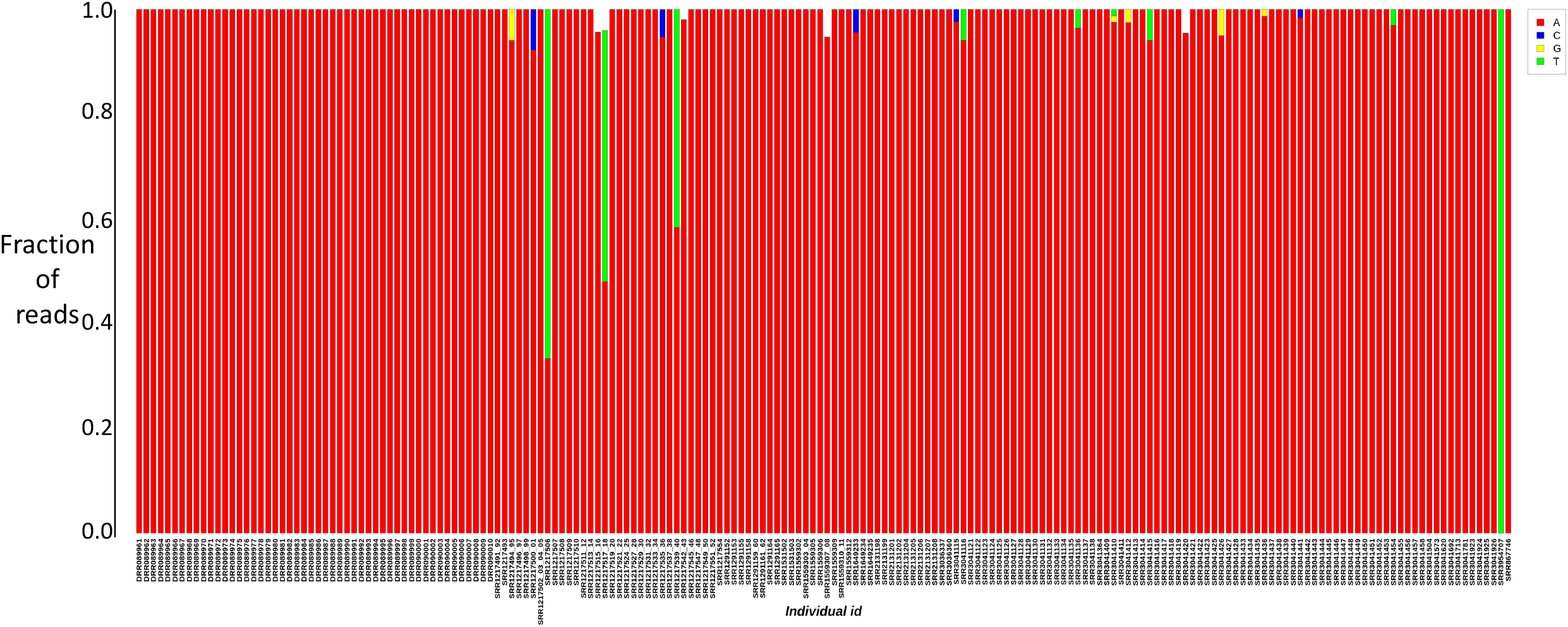
Fraction of reads supporting each of the four nucleotide bases at the premature stop codon inducing site in the *CYP8B1* gene of chicken. Each bar in the figure represents one individual (Short Read Archive ID provided along the x-axis) with the colour red representing the base A and green representing the stop codon inducing base T.

### Candidate species whose bile composition needs further investigation

Based on our analysis of the *CYP8B1* gene across more than 200 species we have identified distinctive and strong patterns of relaxed selection in 14 new species (5 birds, 2 bat species, 3 rodents, 2 primates and one member each of the orders *Eulipotyphla* and *Diprotodontia*) consisting of *Tinamus guttatus, Nipponia nippon, Buceros rhinoceros silvestris, Chaetura pelagica, Anas platyrhynchos, Pteropus vampyrus, Hipposideros armiger, Mus pahari, Fukomys damarensis, Castor canadensis, Eulemur macaco, Carlito syrichta, Sorex araneus* and *Vombatus ursinus*. The distinctive bile composition reported earlier in *Palaeognathae*, hornbills, mousebird and duck seems to be associated with signatures of relaxed selection that we detect in these birds (Hofmann et al. 2010). Although we list 14 species as being under relaxed selection, only the signature found in the *Vombatus ursinus* (apart from the previously reported *Loxodonta africana*) is significant after multiple testing corrections. High quality gap-free genomes for these species might help identify gene loss events in genes of the bile pathway. The presence of a complete open reading frame and lack of clear signals of relaxed selection in the rock hyrax (*Procavia capensis*) suggests that the previously reported bile composition for this species might need to be re-evaluated. However, we cannot rule out the possibility that lack of cholic acid is not reflected in relaxed selection on the *CYP8B1* gene of the rock hyrax.

## Discussion

While comprehensive genome-wide studies that rely upon the abundance of publically available genome sequence data have improved our understanding of gene gain/loss dynamics during evolution of various clades (Hahn et al. 2007b, a), the quality of genomes used and the stringency of quality controls implemented in different studies have remained points of concern despite the maturity of the field (Han et al. 2013; Bornelöv et al. 2017). Gene loss events have generally been inferred based on one of the following types of observations, (a) Gene sequence is completely missing in the region syntenic to the ortholog and no traces of its existence in the form of pseudogenes or partial gene fragments can be found (b) Partial scattered fragments of the gene can be found but are lacking a reading frame (c) The erstwhile open reading frame can be seen to contain multiple stop codon insertions/reading frame disrupting changes (d) The open reading frame has acquired a single pre-mature stop codon and (e) The open reading frame is intact, but the gene is not transcribed or properly spliced or translated due to change or loss of regulatory sequences. Observations of type (a) and (b) can be the result of gaps in the genome assembly or errors during the assembly process and can generally be remedied by searching for the gene sequence in the high coverage raw sequencing read datasets. The best form of evidence for a gene loss event is probably an observation of type (c) in which the open reading frame can still be discerned but has accumulated multiple stop codon insertions and other reading frame disrupting mutations. However, it has to be noted that genes which have accumulated multiple coding frame disrupting changes can take on alternative roles such as regulatory non-coding RNA (Groen et al. 2014). Inferring gene loss from observations of type (d) is probably the most difficult, as the phenotypic consequences of such loss-of-function mutations are still being understood (Sulem et al. 2015; Pagel et al. 2017). Similarly, inference of gene loss based on splice-site disrupting changes and modifications to the gene or splice-site regulatory regions generally require supporting evidence from other functional data.

Use of genomes of single individuals of a species for inference of selection (Mugal et al. 2014) as well as demography (Vijay et al. 2018) have been shown to result in an incomplete picture. Hence, it is not surprising that gene loss inferences are also prone to be affected by limited population level sampling. Yet, comparative genomics across hundreds of species rely on pre-existing datasets and in most cases are able to sample the genome of utmost one individual per species (Hiller et al. 2012; Sharma et al. 2018). However, a clever workaround has been the use of tests of relaxed selection in the focal species compared to a set of background species in a hypothesis testing framework as evidence in support of gene loss (Wertheim et al. 2015; Hecker et al. 2017). While tests of relaxed selection are becoming essential when population genetic sampling is sparse or non-existent, use of such tests requires a sufficiently large number of background species in which this particular gene is still under purifying selection. Moreover, relaxed selection need not necessarily lead to loss of a gene even if it accumulates a stop codon inducing polymorphism. Similarly, it is possible that a species which has acquired a gene loss polymorphism lacks signatures of relaxed selection due to existence of balancing/diversifying selection acting on the gene within the focal species.

In contrast to this, sequencing of thousands of individuals from diverse human populations has allowed the reliable and robust estimation of allele frequencies across the human genome. Efforts are on to assemble Pan-genomes that incorporate population level variation into genome assemblies to better represent the genetic variation found within a species (Sherman et al. 2019). Intriguingly, some of the single nucleotide polymorphisms identified from such large-scale population sampling result in gain/loss of stop codons (MacArthur and Tyler-Smith 2010; MacArthur et al. 2012). In fact, loss of function (LoF) sequence polymorphisms have been frequently observed when allele frequency estimates are based on sampling of large numbers of individuals (Lee and Reinhardt 2012). It has also been argued that LoF variants could have no observable impact on the phenotype (Pagel et al. 2017) and could be rescued by translational stop codon readthrough that has been observed in numerous genes in metazoan species (Jungreis et al. 2011; Dunn et al. 2013; Loughran et al. 2014). Comparative genomic approaches that integrate ribosomal profiling data and proteomics have even suggested a regulatory role for such stop codon readthrough events (Jungreis et al. 2016). Whether some of these stop codons might have some biological or evolutionary significance is still being investigated (Potapova et al. 2018). Hence, gene loss inferences based on single loss of function substitutions have to be treated with caution. We have focussed on the population gene dispensability of the *CYP8B1* gene in chicken in the current study to highlight this issue. Similar screening of population genomic data across other species could potentially identify segregating gene disrupting nucleotide changes. However, this will require availability of genome re-sequencing data from multiple individuals of each species organised as easily accessible pan-genomes.

Sharma & Hiller show that the *CYP8B1* gene in naked mole rat is completely missing in the region syntenic to the human ortholog and represents gene loss event observation of the type (a) defined above. The elephant ortholog of *CYP8B1* has retained the erstwhile open reading frame, but has accumulated four premature stop codons and represents gene loss event observation of the type (c). However, in the case of the manatee, only a single premature stop codon has accumulated in the gene and results in a 35% reduction in the amino acid sequence length. This gene loss inference would be an observation of type (d). Since, gene loss events of type (d) are difficult to understand, Sharma & Hiller rely upon the existence of signatures of relaxed purifying selection in the elephant and manatee lineages as additional evidence in support of the gene loss event. In contrast to the recurrent gene loss pattern identified by Sharma & Hiller it is possible for genes to persist despite relaxed selection or even modify their function (Lahti et al. 2009). Phenotypic plasticity has also been shown to result from relaxed selection (Hunt et al. 2011). However, phenotypic plasticity mediated by changes at the gene expression level that are restricted to specific life stages are harder to identify than changes in the genomic sequence. Hence, being able to identify changes in the coding sequences that reflect changes in regulatory regions would be extremely useful.

The accumulation of multiple stop codon/frame-shift changes in a gene is generally considered strong evidence for the loss of that gene as the truncated protein is unlikely to perform the role of the full protein. Detailed quantification of the functional changes at the phenotypic level is required to ascertain the significance of premature stop codons. Another proxy for identifying changes in function is to look for signatures of intense relaxed selection. Our study uses the *CYP8B1* gene sequence of more than 200 species to look for signatures of relaxed selection across bird and mammal species. Notably, we find that reductions in the degree of purifying selection were detected in lineages that don’t have coding frame disrupting substitutions in the *CYP8B1* gene and these species might have potentially lost other genes in the bile pathway (see **Supplementary Table S4**). Similarly, the prevalence of the stop codon polymorphism in *CYP8B1* gene of chicken can be better understood by looking at different domesticated chicken breeds; its functional consequences on cholesterol metabolism and bile salt composition. Overall, we find that shifts in diet play an important role in determining the selection landscape of *CYP8B1* and other bile pathway genes.

## Acknowledgement

NV would like to acknowledge funding from IISER Bhopal under grant# INST/BIO/2017/019. We thank Council of Scientific & Industrial Research for fellowship to SSS, Ministry of Human Resource Development for fellowships to LT and SS. NV has been awarded the Innovative Young Biotechnologist Award 2018 from the Department of Biotechnology and Early Career Research Award from the Department of Science and Technology (both Government of India).

## Notes

https://github.com/ceglab/CYP8B1

