## Supplementary Text for "Signatures of relaxed selection in the *CYP8B1* gene of birds and mammals"

‘

1. **Genome assembly correction of the *CYP8B1* gene sequence:**
a. **Polar bear**

The polar bear (*Ursus maritimus*) genome has annotated the *CYP8B1* gene by introducing two gaps to correct the reading frame. However, careful comparison of this gap region with other mammals shows that it is missing approximately 38 base pairs. To verify whether this is an assembly artefact, we searched the raw sequencing read data from the individual used for assembling the polar bear genome. We found that the read SRA: SRR933763.27282155.2 spanned across the two base-pair gap and contained a longer stretch of sequence that was missing in the polar bear genome. The genome assembly was corrected by replacing “NNA” with the sequence “ATTGTGGCCCTCTTTCCCTATCTGTCAGTGCAC”. The corrected gene sequence matched the gene sequence of other species from the order *Carnivora* (*Canis familiaris*, *Odobenus rosmarus divergens*, *Leptonychotes weddellii*, *Ailurus fulgens styani*, *Neovison vison*). The same gene sequence was verified in the resequencing data of *Ursus arctos* individuals (see **Supplementary Table S6**). Our curation of Ursus genomes is of special relevance as these species are known to have distinctive bile compositions. We reconstructed the *CYP8B1* gene sequence of the Asiatic black bear (*Ursus thibetanus*) using the polar bear sequence as the reference based on re-sequencing data from the short-read archive. The complete open reading frame could be reconstructed using this mapping approach. The bile composition of the Asiatic black bear is reported to consist only of CDCA and UDCA. In addition to the North American beaver (*Castor canadensis*), the Asiatic black bear is the only other species that is reported to have bile composition of only CDCA and UDCA (Hagey et al. 2010b). The presence of the complete open reading frame in both species despite the signatures of relaxed selection suggest that the *CYP8B1* gene still plays a role in species that have CDCA and UDCA as the major bile components.

b. **Little brown bat**

The complete open reading frame of the *CYP8B1* gene is annotated in ten bat species (*Pteropus vampyrus, Pteropus alecto, Rousettus aegyptiacus, Hipposideros armiger, Desmodus rotundus, Rhinolophus sinicus, Eptesicus fuscus, Myotis brandtii, Myotis davidii, Miniopterus natalensis*). However, in the little brown bat (*Myotis lucifugus*), the *CYP8B1* locus has been annotated with a pseudogene (ENSMLUG00000016464). A premature stop codon that leads to truncations of 85% of the protein is present in the Myoluc2.0 genome assembly. We investigated the support for this premature stop codon in the raw sequencing read data and found that this was actually an assembly error potentially caused due to a homo-polymer run (six G’s assembled instead of five G’s supported by raw reads). RNA-seq data from six different *Myotis lucifugus* individuals supporting the correction of the genome assembly had multiple reads from each individual (see **Supplementary Table S6**). Our analysis suggests that the genome assembly of the little brown bat needs to be corrected at this position. High coverage whole genome sequencing data will need to be used to corroborate this result.

c. **Canary**

The current assembly of *Serinus canaria* requires a one base pair gap to maintain the open reading frame of the *CYP8B1* gene sequence. Search of the raw sequencing read data showed that the sequence around the gap represented by an N should be corrected from “CTG**N**CTGAC” to “CTGCT***A***GAC”/ “CTGCT***G***GAC”. The sixth site in this sequence is polymorphic (A/G) and might have caused the assembly error. List of reads supporting this correction are provided in **Supplementary Table S6**.

d. **Platypus**

The Platypus (*Ornithorhynchus anatinus*) genome annotation has a longer C-terminal tail in the *CYP8B1* gene. However, careful analysis of read support showed that the assembly had erroneously included an N instead of a T. The files with reads supporting the corrected sequence have been provided as SAM files (see <https://github.com/ceglab/CYP8B1>**/SAMs**).

**1.2 Gene sequence verification using raw sequencing reads**

In order to ascertain the correctness of the gene sequences present in the NCBI and Ensembl databases we verified the *CYP8B1* gene sequences against raw sequencing reads from the Short Read Archive. The *CYP8B1* complete open reading frame was searched against short read archive datasets of the same species using SRA blastn utility. The resulting blast hits were saved as SAM files (see <https://github.com/ceglab/CYP8B1>**/SAMs**). Read alignments were visualised using IGV browser to manually verify the gene sequences.

2. **Re-annotation of the *CYP8B1* gene:**

The *CYP8B1* gene is annotated in the human genome as a single exon gene that produces a protein of length 501 amino acids. However, in some species the *CYP8B1* gene is annotated by ensembl 94 to consist of more than one exon. Moreover, in some species the annotated gene does not contain the complete open reading frame. Hence, we manually curated the annotation of this gene prior to using it for multiple sequence alignment and downstream analysis. DNA ambiguity characters were randomly replaced by one of the possible DNA bases. The re-annotated open reading frames and their multiple sequence alignment are provided as <https://github.com/ceglab/CYP8B1>**/ORFs** and <https://github.com/ceglab/CYP8B1>**/MSAs**.

The annotation for the ferret (*Mustela putorius furo*) *CYP8B1* gene was corrected by extending the coding region annotation by three codons till the stop codon to reach a protein of length 501 amino acids. The transcript annotated in the current ensembl build (version 94) in the cow (*Bos taurus*) genome consists of two exons that results in a protein of length 166 amino acids. We extracted the genomic nucleotide sequence stretching from the beginning of the first exon till the end of second exon and searched for open reading frames. The longest continuous open reading frame was found to begin at the previously annotated start codon and extended past the boundary of the first exon to produce an open reading frame coding for a protein of length 499 amino acids. We believe this is a more parsimonious annotation as it consists of fewer exons and produces a longer protein; however, the existence of an additional exon that codes for an alternative splice isoform would need to be evaluated.

The goat genome assembly is of an extremely high quality, leading to it being considered a benchmark for future genome assembly projects (aka, the golden goat) (Worley 2017). Since the goat *CYP8B1* gene has two exons annotated in ensembl 94 we reasoned that the high quality goat genome might have correctly assembled and annotated the gene. However, we found that the additional goat specific exon had a stretch of only 10 codons before reaching the stop codon. The longest continuous ORF that extended past the end of exon1 into the first intron resulted in a gene of 499 amino acids compared to the 498 amino acid long gene that has currently been annotated in ensembl 94. Our corrected annotation of the goat *CYP8B1* gene was extremely similar to the gene sequence in the sheep genome. Although we cannot rule out the possibility of an alternative isoform, the single exon annotation is better supported across other species annotation. The rationale and methodology used to re-annotate the *CYP8B1* gene in the cow and goat genome was used for other species that had multi-exon annotations for the *CYP8B1* gene (see **Supplementary Table S7**). Overall, the *CYP8B1* gene was re-annotated in 18 genomes to obtain a gene sequence that is comparable to the human genome annotation. Our re-annotated open reading frames are supported by similar annotation by the NCBI automated annotation pipeline.

3. **Analysis of population genetic variation across chicken breeds:**

We downloaded the re-sequencing data from multiple chicken breeds from the Short Read Archive (SRA) and mapped it to the chicken genome using the bwa mem read mapper (Li 2013). Details of the datasets downloaded and associated breed details are provided in **Supplementary Table S10**. The number of reads supporting each of the bases at a given position in the read alignments was calculated using the bam read-count utility (github.com/genome/bam-readcount). These read-count values were further verified by manual inspection using the IGV (Integrative Genomics Viewer) browser (Robinson et al. 2011).

4. **Lineage specific analysis:**
a. **Cetartiodactyla**

The sixteen cetacean species included in our analysis consists of *Lipotes vexillifer*, *Sousa chinensis*, *Sousa sahulensis*, *Tursiops truncatus*, *Tursiops aduncus*, *Orcinus orca*, *Lagenorhynchus obliquidens, Delphinapterus leucas*, *Phocoena phocoena*, *Neophocaena asiaeorientalis asiaeorientalis, Physeter catodon*, *Balaenoptera acutorostrata scammoni*, *Balaenoptera bonaerensis*, *Balaenoptera physalus*, *Eschrichtius robustus* and *Balaena mysticetus*. Artiodactyla species included the hippopotamus (*Hippopotamus amphibius*), giraffe (*Giraffa Camelopardalis*), pig, camels, sheep, goat, cattle, alpaca (*Vicugna pacos*), antelope (*Pantholops hodgsonii*) and deer (*Elaphurus davidianus*, *Cervus elaphus hippelaphus*, *Capreolus capreolus* and *Odocoileus virginianus*). We verified the validity of the shorter open reading frame in the hippopotamus (*Hippopotamus amphibius*) and the longer open reading frame in *Orcinus orca* and *Lagenorhynchus obliquidens* using short read archive data (see <https://github.com/ceglab/CYP8B1>**/SAMs**).

b. **Afrotheria**

Other than the elephant and manatee other species from Afrotheria clade whose genomes have been sequenced include the tenrec, hyrax, aardvark, cape elephant shrew and golden mole species. The lesser hedgehog tenrec (*Echinops telfairi*) genome assembly EchTel2.0 has been annotated with a 501 amino acid long open reading frame for the *CYP8B1* gene. Similarly, the aardvark (*Orycteropus afer afer*: OryAfe1.0), cape elephant shrew (*Elephantulus edwardii*: EleEdw1.0) and golden mole (*Chrysochloris asiatica*: ChrAsi1.0) genomes also have full length open reading frames of 501 amino acids each. The existence of a full length ORF in the representative members of *Tenrecoidea* (i.e., *Echinops telfairi*), *Chrysochloridae* (i.e., *Chrysochloris asiatica*), *Tubulidentata* (i.e., *Orycteropus afer afer*) and *Macroscelidea* (i.e., *Elephantulus edwardii*) suggests that the *CYP8B1* gene is probably present in other species of *Afroinsectivora*. Although the streaked tenrec (*Hemicentetes semispinosus*) bile is reported to have C_27_ alcohols, it also contains C_24_ 5β-acids and would be an interesting species to investigate when genetic data is available (Hagey et al. 2010b). Despite having a complete ORF in the cape golden mole (*Chrysochloris asiatica*), the *CYP8B1* gene is known to be under relaxed selection. The bile pathway is further compromised in the cape golden mole by the loss of the *SLC27A5* gene (Sharma and Hiller 2018). It has been experimentally demonstrated that the *CYP8B1* gene is regulated by Hepatocyte nuclear factor 4-alpha (HNF4A) (Zhang and Chiang 2001). The transcription start site and HNF4A binding site have been delineated and subsequently validated using the CHIP-seq approach in human, mouse and rat (Schmidt et al. 2010). Despite the presence of coding frame disrupting mutations in the *CYP8B1* gene of the elephant the regulatory region is still conserved and the gene continues to get transcribed. Multiple reads from the gene can be found in the RNA-seq data of the African elephant.

We could annotate a complete ORF of 500 amino acids in the latest genome assembly of the cape elephant shrew (*Procavia capensis*: Pcap_2.0). The presence of a complete open reading frame in the rock hyrax is a bit unexpected as it has been grouped along with the elephants and manatees in a previous study that focussed on bile salt composition (Hagey et al. 2010b). However, the raw sequencing read data for the rock hyrax (*Procavia capensis*) supported the existence of a complete ORF. The existence of the *CYP8B1* gene in the representative member of *Hyracoidea* (i.e., *Procavia capensis*) hints at the loss of the gene within *Tethytheria* (*Proboscidea* and *Sirenia*). Sequencing of more genomes within afrotherian lineages will help further delineate the timing of this gene loss event. Understanding the timing of this gene loss event may help us understand the reason for relaxed selection in the *CYP8B1* gene and its subsequent loss.

c. **Perissodactyla and carnivores**

Within Perissodactyla, equids and tapirs are reported to have cholic acid in their bile. However, all rhinoceros species are thought to have a bile profile similar to elephants and manatee. Despite the lack of cholic acid, both the Sumatran and southern white rhino genomes seem to retain a complete open reading frame for the *CYP8B1* gene. Signatures of relaxed selection have previously been reported in the southern white rhino (Sharma and Hiller 2018). We looked for relaxed selection within perissodactyla using sequences of both the Southern white rhino and Sumatran rhino (*Dicerorhinus sumatrensis*) along with *Equus caballus*, *Equus asinus* and *Equus przewalskii*. Strong relaxed selection was consistently identified in the Sumatran rhino. While relaxed selection could be seen in the white rhino, it was not significant when the Sumatran rhino was included in the background set. However, relaxed selection in the white rhino was stronger when the Sumatran rhino was excluded. Strong intensification of selection can be seen in *Equus asinus*.

None of the carnivore species analysed in our study showed significant relaxed selection. However, the general descriptive model on the alignment of carnivores shows strong signatures of relaxed selection in *Neomonachus schauinslandi*, *Odobenus rosmarus divergens* and *Arctocephalus gazelle*. The grizzly bear (*Ursus arctos horribilis*), the polar bear (*Ursus maritimus*), the Asiatic black bear (*Ursus thibetanus*) and the black bear (*Ursus americanus*) are known to produce ursodeoxycholic acid (Hofmann et al. 2010). Nonetheless, cholic acid is also reported in the bile of bear species except for the Asiatic black bear (*Ursus thibetanus*). Consistent with this we see strong relaxation of selection in the Asiatic black bear lineage (**Supplementary Figure S3C**). However, relaxed selection is seen in the black bear (*Ursus americanus*) and grizzly bear (*Ursus arctos horribilis*) lineages as well. Surprisingly, the polar bear (*Ursus maritimus*) lineage shows intensification of selection. These results are in line with a recent study that found changes in the amylase gene content of the polar bear that represents a transition from omnivorous to a mostly carnivorous diet (Rinker et al. 2019). Fine-scale quantification of the relative amounts of the various bile components across bears might help better understand these patterns.

d. **Rodents and Primates**

The rodent species with completely sequenced genomes that are evolutionarily closely related to the Naked-mole rat include the Damara mole rat (*Fukomys damarensis*), Long-tailed chinchilla (*Chinchilla lanigera*) and Common degu (*Octodon degus*). All three of these species have complete open reading frames of the *CYP8B1* gene. In addition to these three species we included sixteen other rodent species in our multi-species alignment. The *CYP8B1* gene of the North American beaver (*Castor Canadensis*) assembled in the latest version of the genome was found to contain two separate single base-pair deletions compared to other rodent species. The illumina reads supporting this version of the *CYP8B1* gene consisted of three different haplotypes, each with a different single base-pair insertion (see <https://github.com/ceglab/CYP8B1>**/SAMs**). Hence, it was unclear whether the gene sequence is the result of an assembly error or repeated segmental duplications. However, an earlier version of the beaver genome did not have the single base-pair deletions seen in the current version of the genome. Hence, we corrected the gene sequence by including the two bases from the older version of the assembly that have been deleted in the latest version of the assembly. The complete open reading frame of the gene could be annotated with this corrected version of the assembly. The reads supporting this corrected version of the gene did not have multiple insertion-containing haplotypes that could be seen with the pre-correction version (see <https://github.com/ceglab/CYP8B1>**/SAMs**).

We found that the *CYP8B1* gene is annotated with a segmentally duplicated paralog in the Domestic guinea pig (*Cavia porcellus*) and the Brazilian guinea pig (*Cavia aperea*). These two copies of the *CYP8B1* gene are located approximately 6Kb apart. One of the copies is exactly 501 amino acids long while the other copy is 558 amino acids long. Although we could not rule out the possibility of a genome assembly error that resulted in this annotation, RNA-seq expression measurements from both wild and domesticated guinea pigs shows evidence for expression of the 501 amino acids long copy but not for the 558 amino acids long duplicate (Albert et al. 2012). Due to ambiguity in genome assembly quality we did not include the gene sequences from either guinea pig species. Detailed investigation of this region with long read sequencing technologies might help establish whether this is a genome assembly error or a true segmental duplication event. None of the 21 primate species included in our study showed significant relaxation of selection. However, intensification of selection was seen in the green monkey (*Chlorocebus sabaeus*) and the drill (*Mandrillus leucophaeus*).

e. **Birds**

The gene sequences of all the birds included in our alignment had a complete open reading frame after the correction of the *Serinus canaria* gene sequence. One stretch of nine amino acids and another of three amino acids was found across all the bird sequences but was missing in the gene sequence of the other species. It is possible that these regions of the *CYP8B1* gene have a bird specific function. However, this would require detailed functional characterisation of these regions. Moreover, the gene order in the vicinity of the *CYP8B1* gene in birds is different from that found in mammals. The *CYP8B1* and the adjacent *ACKR2* gene are positioned along the chromosome in the same orientation in birds but in opposing directions in mammals. The other adjacent gene is *FAM198A* in mammals but in birds it is *OBSCN*. Bird species that showed strong relaxation or intensification are discussed along with what is known about their bile composition.

a. **Palaeognathae**

Previous studies that have profiled the bile composition of palaeognathae species have found C_27_ alcohols and C_27_ acids within Tinamidae and C_27_ acids in *Struthiformes* (Hagey et al. 2009, 2010a). Although the *CYP8B1* gene sequences are not available for the exact same species of tinamou, we found relaxed selection in the white-throated tinamou (*Tinamus guttatus*). Strong relaxed selection can also be seen in the emu (*Dromaius novaehollandiae*), but does not qualify the significance criteria (**Supplementary Table S4**).

b. **Passeriformes**

None of the 22 Passeriformes species present in our alignment showed consistent relaxed selection (**Supplementary Table S4**). However, we found intensified selection in the Javan myna (*Acridotheres javanicus*), Lawe’s parotia (*Parotia lawesii*) and the barn swallow (*Hirundo rustica rustica*).

c. **Telluraves** *(without passerines)*

Among the 14 species from the clade Telluraves *(without passerines)*, we find evidence for relaxed selection (**Supplementary Table S4**) in the rhinoceros hornbill (*Buceros rhinoceros silvestris*) and the speckled mousebird (*Colius striatus*). The rhinoceros hornbill (*Buceros rhinoceros*) has a bile composition consisting of a mixture of C_27_ bile acids and C_27_ bile alcohol sulfates. The speckled mousebird is known to have a bile composition consisting of a mixture of C_24_ bile acids, C_27_ bile acids and C_27_ bile alcohol sulfates (Hagey et al. 2010a).

d. **Aequorlitornithes**

Among the 16 species from the clade *Aequorlitornithes*, we find evidence for relaxed selection (**Supplementary Table S4**) in the crested ibis (*Nipponia nippon*) and intensified selection in the killdeer (*Charadrius vociferus*).

e. **Ducks**

We looked for signatures of relaxed selection in 4 species of the order Anseriformes and found evidence (**Supplementary Table S4**) of relaxed selection in the mallard (*Anas platyrhynchos*). It would be interesting to investigate whether the signature of relaxed selection identified here are reflective of regulatory changes that might play a role in Duck lipid traits (Zhang et al. 2018). Although, we could find a large gap in the ensemble alignment of the duck genome with other bird species in the upstream region of the *CYP8B1* gene, it is unclear at this stage whether this gap corresponds to an actual change in the regulatory sequence of the *CYP8B1* gene of the duck.

**Supplementary tables**

**Supplementary Table S1**: Consistency of multiple sequence alignments of the *CYP8B1* gene calculated using the program SuiteMSA.

| **Alignment pair** | **% consistency** | **Sum of pairs score** | **Column score** | **Column score**  **(with gaps)** |
| --- | --- | --- | --- | --- |
| CLUSTALW & MAFFT | 78.311 | 0.989 | 0.97 | 0.796 |
| CLUSTALW & MUSCLE | 76.549 | 0.989 | 0.947 | 0.776 |
| CLUSTALW & PRANK | 69.474 | 0.986 | 0.957 | 0.768 |
| MAFFT & MUSCLE | 77.219 | 0.999 | 0.951 | 0.779 |
| MAFFT & PRANK | 72.331 | 0.997 | 0.968 | 0.787 |
| MUSCLE & PRANK | 68.722 | 0.996 | 0.959 | 0.761 |

**Supplementary Table S2**: Comparison of multiple sequence alignments of the *CYP8B1* gene from four different programs by MUMSA (Lassmann and Sonnhammer 2005) in terms of AOS (Average Overlap Score) and MOS (Multiple Overlap Score). The nucleotide substitution model selected by modeltest-ng as the best model based on BIC is also listed.

| **Sl. No** | **Species group** | **AOS** | **MOS**  (PRANK) | **MOS**  (MUSCLE) | **MOS**  (CLUSTALW) | **MOS**  (MAFFT) | **Substitution**  **model selected** |
| --- | --- | --- | --- | --- | --- | --- | --- |
| 1 | Aequorlitornithes | 0.75 | 0.99 | 0.99 | 0.99 | 0.99 | HKY+G4 |
| 2 | *Afrotheria* | 0.75 | 1.0 | 1.0 | 1.0 | 1.0 | HKY+G4 |
| 3 | *Anseriformes* |  |  |  |  |  |  |
| 4 | *Chiroptera* | 0.75 | 0.99 | 0.99 | 0.99 | 0.99 | HKY+G4 |
| 5 | *Carnivora* | 0.75 | 1.0 | 1.0 | 1.0 | 1.0 | HKY+G4 |
| 6 | *Cetacean* | 0.75 | 1.0 | 1.0 | 1.0 | 1.0 | HKY+G4 |
| 7 | Columbaves, Strisores & Gruiformes | 0.75 | 1.0 | 1.0 | 1.0 | 1.0 | HKY+G4 |
| 8 | Galliformes | 0.75 | 1.0 | 1.0 | 1.0 | 1.0 | HKY+G4 |
| 9 | Lemur | 0.75 | 0.99 | 0.99 | 0.99 | 0.99 | HKY+I |
| 10 | Marsupial | 0.72 | 0.96 | 0.97 | 0.96 | 0.97 | GTR+I |
| 11 | Paleognathae | 0.75 | 1.0 | 1.0 | 1.0 | 1.0 | HKY+G4 |
| 12 | Passeriformes | 0.75 | 0.99 | 0.99 | 0.99 | 0.99 | HKY+G4 |
| 13 | Primates | 0.75 | 1.0 | 1.0 | 1.0 | 1.0 | HKY+G4 |
| 14 | Perissodactyla | 0.75 | 1.0 | 1.0 | 1.0 | 1.0 | HKY |
| 15 | Rodents | 0.75 | 0.99 | 0.99 | 0.99 | 0.99 | HKY+I+G4 |
| 16 | Telluraves | 0.75 | 0.99 | 0.99 | 0.99 | 0.99 | HKY+G4 |

**Supplementary Table S3**: Results of sequence substitution saturation tests described in (Xia et al. 2003) and implemented in the program DAMBE (Xia 2013). The index to measure substitution saturation (Iss) and the corresponding critical Iss (Iss.c) estimates for symmetrical and extreme asymmetrical tree are reported for each group of species that has been used in our study.

| **Sl. No** | **Species group** | **Iss** | **Iss.c**  (symmetrical) | **Iss.c**  (asymmetrical) |
| --- | --- | --- | --- | --- |
| 1 | Aequorlitornithes | 0.088 | 0.7929 | 0.6098 |
| 2 | Afrotheria | 0.1034 | 0.7847 | 0.5675 |
| 3 | Carnivores | 0.1034 | 0.7847 | 0.5675 |
| 4 | Cetaceans (4 OTUs) | 0.075 | 0.833 | 0.803 |
| 5 | Cetaceans (8 OTUs) | 0.074 | 0.809 | 0.708 |
| 6 | Cetaceans (16 OTUs) | 0.076 | 0.792 | 0.608 |
| 7 | Cetaceans (32 OTUs) | 0.078 | 0.774 | 0.49 |
| 8 | Chiroptera | 0.1209 | 0.7905 | 0.6532 |
| 9 | Columbaves Strisores & Gruiformes | 0.1242 | 0.7978 | 0.6855 |
| 10 | Galliformes | 0.082 | 0.812 | 0.745 |
| 11 | Lemur | 0.1926 | 0.8112 | 0.7439 |
| 12 | Marsupial | 0.4216 | 0.8105 | 0.7429 |
| 13 | Paleognathae_with_tortoise | 0.166 | 0.8063 | 0.7228 |
| 14 | Passeriformes | 0.0977 | 0.7849 | 0.5688 |
| 15 | Perissodactyla | 0.0471 | 0.8178 | 0.7684 |
| 16 | Primate | 0.0426 | 0.7847 | 0.5727 |
| 17 | Rodents | 0.2447 | 0.784 | 0.5825 |
| 18 | Telluraves | 0.124 | 0.7863 | 0.6203 |

**Supplementary Table S4**: Results of performing tests for relaxed selection in each of the species included in our multiple sequence alignment using the RELAX program and test for episodic diversifying selection using aBSREL. The K-value represents the intensity of selection. Significant result in the hypothesis test with values of K < 1 refer to relaxed purifying selection and K>1 refers to intensification of selection.

| **Test Species** | **p-value** | | **K** | | **Relaxation**  (after FDR correction  q-value < 0.1) | **Diversifying positive selection**  **(p-value)** |
| --- | --- | --- | --- | --- | --- | --- |
| ***Cetartiodactyla*** | | | | | | |
| *Lipotes vexillifer* | 0.1159 | 0.25 | | | Not significant | NA |
| *Delphinapterus leucas* | 0.495 | 8.98 | | | Not significant | NA |
| *Neophocaena asiaeorientalis asiaeorientalis* | 0.2683 | 50 | | | Not significant | NA |
| *Phocoena phocoena* | 0.2594 | 50 | | | Not significant | NA |
| *Lagenorhynchus obliquidens* | 0.2995 | 0 | | | Not significant | NA |
| *Sousa sahulensis* | 0.3195 | 0 | | | Not significant | NA |
| *Sousa chinensis* | 0.8969 | 0.73 | | | Not significant | NA |
| *Orcinus orca* | 0.448 | 0.07 | | | Not significant | NA |
| *Tursiops aduncus* | 0.4189 | 0.16 | | | Not significant | NA |
| *Tursiops truncatus* | 0.8511 | 0.75 | | | Not significant | NA |
| *Vicugna pacos* | 0.752 | 1.09 | | | Not significant | NA |
| *Physeter catodon* | 0.1797 | 0.35 | | | Not significant | NA |
| *Capra sibirica* | 0.0626 | 50 | | | Not significant | NA |
| *Capra hircus* | 0.7341 | 0.46 | | | Not significant | NA |
| *Ammotragus lervia* | 0.1292 | 50 | | | Not significant | NA |
| *Ovis aries* | 0.7166 | 0.57 | | | Not significant | NA |
| *Pantholops hodgsonii* | 0.703 | 2.66 | | | Not significant | NA |
| *Bos indicus* | 0.1262 | 50 | | | Not significant | NA |
| *Bos taurus* | 0.4447 | 0.06 | | | Not significant | NA |
| *Bison bison bison* | 0.8667 | 0.95 | | | Not significant | NA |
| *Bos mutus* | 0.687 | 0.5 | | | Not significant | NA |
| *Camelus bactrianus* | 0.6611 | 5.14 | | | Not significant | NA |
| *Bubalus bubalis* | 0.7544 | 2.52 | | | Not significant | NA |
| *Giraffa camelopardalis tippelskirchi* | 0.0638 | 7.73 | | | Not significant | NA |
| *Capreolus capreolus* | 0.0818 | 8.69 | | | Not significant | NA |
| *Cervus elaphus hippelaphus* | 0.5107 | 4.95 | | | Not significant | NA |
| *Elaphurus davidianus* | 0.72 | 2.26 | | | Not significant | NA |
| *Odocoileus virginianus texanus* | 0.1279 | 14.77 | | | Not significant | NA |
| *Sus scrofa* | 0.3359 | 0.43 | | | Not significant | NA |
| *Condylura cristata* | 0.0539 | 0 | | | Not significant | NA |
| *Camelus dromedarius* | 0.8336 | 0.59 | | | Not significant | NA |
| *Hippopotamus amphibius* | 0.7972 | 1.23 | | | Not significant | NA |
| *Balaena mysticetus* | 0.4057 | 0.25 | | | Not significant | NA |
| *Eschrichtius robustus* | 0.7277 | 0.53 | | | Not significant | NA |
| *Balaenoptera bonaerensis* | 1 | 0.28 | | | Highly unreliable K | NA |
| *Balaenoptera acutorostrata scammoni* | 0.429 | 0.18 | | | Not significant | NA |
| *Balaenoptera physalus* | 0.2483 | 13.27 | | | Not significant | NA |
| ***Afrotheria*** | | | | | | |
| *Elephantulus edwardii* | 0.0151 | 7.38 | | Not significant | | NA |
| *Chrysochloris asiatica* | 0.1882 | 0.73 | | Not significant | | NA |
| *Orycteropus afer afer* | 0.2628 | 0.32 | | Not significant | | NA |
| *Procavia capensis* | 0.3433 | 1.38 | | Not significant | | NA |
| *Loxodonta africana* | 0.0000 | 0.03 | | **Relaxed (gene loss)** | | NA |
| *Trichechus manatus latirostris* | 0.1821 | 0.46 | | Not significant | | NA |
| *Echinops telfairi* | 0.0073 | 1.44 | | Not significant | | 0.01179 |
| ***Chiroptera*** | | | | | | |
| *Pteropus vampyrus* | 0.0009 | | 10.3/0.0 | | Highly unreliable K | NA |
| *Pteropus alecto* | 0.0078 | | 5.38 | | Not significant | NA |
| *Rhinolophus sinicus* | 0.6111 | | 0.16 | | Not significant | NA |
| *Hipposideros armiger* | 0.0113 | | 0.0 | | Highly unreliable K | NA |
| *Myotis davidii* | 0.1642 | | 1.26/0.0 | | Highly unreliable K | NA |
| *Myotis lucifugus* | 0.0975 | | 10.88 | | Not significant | NA |
| *Myotis brandtii* | 0.3941 | | 0.0 | | Not significant | NA |
| *Miniopterus natalensis* | 0.0213 | | 1.72 | | Not significant | NA |
| *Desmodus rotundus* | 0.2487 | | 0.07 | | Not significant | NA |
| *Eptesicus fuscus* | 0.2745 | | 0.0 | | Not significant | NA |
| ***Carnivora*** | | | | | | |
| *Arctocephalus gazelle* | 0.0869 | | 0.00 | | Not significant | NA |
| *Odobenus rosmarus divergens* | 0.1747 | | 0.00 | | Not significant | NA |
| *Neovison vison* | 0.4577 | | 2.44 | | Not significant | NA |
| *Enhydra lutris kenyoni* | 0.4327 | | 0.77 | | Not significant | NA |
| *Ailurus fulgens styani* | 0.7106 | | 1.08 | | Not significant | NA |
| *Puma concolor* | 0.6015 | | 1.31 | | Not significant | NA |
| *Felis catus* | 0.5341 | | 1.33 | | Not significant | NA |
| *Panthera tigris altaica* | 0.5717 | | 1.33 | | Not significant | NA |
| *Panthera pardus* | 0.68 | | 0.71 | | Not significant | NA |
| *Hyaena hyaena* | 0.49 | | 1.15 | | Not significant | NA |
| *Lycaon pictus* | 0.7229 | | 0.69 | | Not significant | NA |
| *Manis javanica* | 0.6526 | | 1.08 | | Not significant | NA |
| *Canis familiaris* | 0.8559 | | 1.21 | | Not significant | NA |
| *Vulpes vulpes* | 0.6467 | | 1.21 | | Not significant | NA |
| *Ailuropoda melanoleuca* | 0.9579 | | 0.98 | | Not significant | NA |
| *Ursus arctos horribilis* | 0.1394 | | 0.0 | | Not significant | NA |
| *Ursus maritimus* | 0.7026 | | 1.29 | | Not significant | NA |
| *Neomonachus schauinslandi* | 0.2013 | | 0.00 | | Not significant | NA |
| *Leptonychotes weddellii* | 0.9579 | | 0.98 | | Not significant | NA |
| *Callorhinus ursinus* | 0.9894 | | 1.06 | | Not significant | NA |
| *Ursus americanus* | 0.4325 | | 0.42 | | Not significant | NA |
| *Ursus thibetanus* | 0.299 | | 0.0 | | Not significant | NA |
| ***Perissodactyla*** | | | | | | |
| *Dicerorhinus sumatrensis sumatrensis* | 0.0253 | | 0.00 | | Not significant | NA |
| *Equus asinus* | 0.0906 | | 43.79 | | Not significant | NA |
| *Equus caballus* | 0.3421 | | 38.44 | | Not significant | NA |
| *Equus przewalskii* | 1.0 | | 0.54/ 1.17 | | Highly unreliable K | NA |
| *Ceratotherium simum simum* | 0.8524 | | 0.88 | | Not significant | NA |
| ***Primates*** | | | | | | |
| *Rhinopithecus roxellana* | 0.9227 | | 1.3 | | Not significant | NA |
| *Rhinopithecus bieti* | 0.3188 | | 36.05/0.00 | | Highly unreliable K | NA |
| *Chlorocebus sabaeus* | 0.0437 | | 2.66 | | Not significant | 0.03155 |
| *Papio anubis* | 0.1654 | | 0.0 | | Not significant | NA |
| *Theropithecus gelada* | 0.8338 | | 0.64 | | Not significant | NA |
| *Cercocebus atys* | 0.8121 | | 1.19 | | Not significant | NA |
| *Mandrillus leucophaeus* | 0.0353 | | 50.0 | | Not significant | NA |
| *Macaca mulatta* | 0.8471 | | 0.63 | | Not significant | NA |
| *Macaca fascicularis* | 1.0 | | 1.16/ 1.0 | | Highly unreliable K | NA |
| *Macaca nemestrina* | 0.8539 | | 1.17 | | Not significant | NA |
| *Cebus capucinus imitator* | 0.7307 | | 1.29 | | Not significant | NA |
| *Aotus nancymaae* | 1.0 | | 1.06/ 1.0 | | Highly unreliable K | NA |
| *Callithrix jacchus* | 0.2651 | | 0.52 | | Not significant | NA |
| *Saimiri boliviensis boliviensis* | 0.1891 | | 0.3 | | Not significant | NA |
| *Gorilla gorilla gorilla* | 0.002 | | 0.00/ 5.62 | | Highly unreliable K | 0.01331 |
| *Pan troglodytes* | 0.8939 | | 1.27 | | Not significant | NA |
| *Pan paniscus* | 0.8754 | | 1.13 | | Not significant | NA |
| *Homo sapiens* | 0.2243 | | 0.0 | | Not significant | NA |
| *Pongo abelii* | 0.8955 | | 0.9 | | Not significant | NA |
| *Colobus angolensis palliates* | 0.8136 | | 0.66 | | Not significant | NA |
| *Piliocolobus tephrosceles* | 0.8339 | | 1.16 | | Not significant | NA |
| ***Prosimians* & *Dermoptera*** | | | | | | |
| *Eulemur macaco* | 0.0025 | | 0.06 | | Not significant | NA |
| *Carlito syrichta* | 0.0169 | | 0.00 | | Not significant | NA |
| *Tupaia belangeri chinensis* | 0.1805 | | 7.21 | | Not significant | NA |
| *Galeopterus variegatus* | 0.638 | | 0.78 | | Not significant | NA |
| *Otolemur garnettii* | 0.2228 | | 0.51 | | Not significant | NA |
| *Propithecus coquereli* | 0.1213 | | 2.25 | | Not significant | 0.03222 |
| ***Monotremata*, *Marsupialia* & *Xenarthra*** | | | | | | |
| *Vombatus ursinus* | 0.0 | | 0.0 | | **Relaxed** | NA |
| *Ornithorhynchus anatinus* | 0.009 | | 7.22 | | Not significant | NA |
| *Erinaceus europaeus* | 0.1493 | | 0.67 | | Not significant | 0.03694 |
| *Dasypus novemcinctus* | 0.1579 | | 0.7 | | Not significant | NA |
| *Sorex araneus* | 0.0209 | | 0.36 | | Not significant | NA |
| *Phascolarctos cinereus* | 0.0 | | 8.44 | | **Intensified** | 0.00338 |
| ***Rodents*** | | | | | | |
| *Peromyscus maniculatus bairdii* | 0.4549 | | 3.9 | | Not significant | NA |
| *Nannospalax galili* | 0.3162 | | 0.64 | | Not significant | NA |
| *Jaculus jaculus* | 0.1416 | | 3.39 | | Not significant | NA |
| *Fukomys damarensis* | 0.0016 | | 0.05 | | Not significant | NA |
| *Castor canadensis* | 0.0313 | | 0.00 | | Not significant | NA |
| *Chinchilla lanigera* | 0.0543 | | 0.04 | | Not significant | NA |
| *Octodon degus* | 0.3209 | | 0.72 | | Not significant | NA |
| *Marmota marmota marmota* | 0.7368 | | 0.72 | | Not significant | NA |
| *Ictidomys tridecemlineatus* | 0.2463 | | 0.15 | | Not significant | NA |
| *Dipodomys ordii* | 0.0017 | | 4.89 | | Not significant | NA |
| *Oryctolagus cuniculus* | 0.037 | | 1.88 | | Not significant | NA |
| *Meriones unguiculatus* | 0.845 | | 0.97 | | Not significant | NA |
| *Rattus norvegicus* | 0.5728 | | 1.37 | | Not significant | NA |
| *Mus pahari* | 0.0122 | | 0.19 | | Not significant | NA |
| *Mus caroli* | 0.0781 | | 3.55 | | Not significant | 0.00352 |
| *Mus musculus* | 1.0 | | 1.19 | | Not significant | NA |
| *Mus spretus* | 0.186 | | 0.04 | | Not significant | NA |
| *Microtus ochrogaster* | 0.1537 | | 0.51 | | Not significant | NA |
| *Cricetulus griseus* | 0.2879 | | 0.82/3.44 | | Highly unreliable K | NA |
| ***Palaeognathae* (without *Gopherus agassizii*)** | | | | | | |
| *Dromaius novaehollandiae* | 0.06 | | 0.33 | | Not significant | NA |
| *Nothoprocta perdicaria* | 0.5571 | | 2.31 | | Not significant | NA |
| *Eudromia elegans* | 0.8856 | | 0.9 | | Not significant | NA |
| *Crypturellus cinnamomeus* | 0.1154 | | 0.0/10.07 | | Highly unreliable K | NA |
| *Tinamus guttatus* | 0.0494 | | 0.28 | | Not significant | NA |
| *Struthio camelus australis* | 0.4555 | | 2.61 | | Not significant | NA |
| ***Palaeognathae* (with *Gopherus agassizii*)** | | | | | | |
| *Dromaius novaehollandiae* | 0.0907 | | 0.38 | | Not significant | NA |
| *Nothoprocta perdicaria* | 0.4552 | | 1.47 | | Not significant | NA |
| *Eudromia elegans* | 0.8364 | | 1.09 | | Not significant | NA |
| *Crypturellus cinnamomeus* | 0.0874 | | 2.25 | | Not significant | NA |
| *Tinamus guttatus* | 0.0973 | | 0.19 | | Not significant | NA |
| *Struthio camelus australis* | 0.5711 | | 1.34 | | Not significant | NA |
| *Gopherus agassizii* | 0.1635 | | 0.67 | | Not significant | NA |
| ***Passeriformes*** | | | | | | |
| *Taeniopygia guttata* | 0.5413 | | 0.89 | | Not significant | NA |
| *Lonchura striata domestica* | 0.5183 | | 0.8 | | Not significant | NA |
| *Serinus canaria* | 0.591 | | 1.15 | | Not significant | NA |
| *Geospiza fortis* | 0.1705 | | 0.53 | | Not significant | NA |
| *Junco hyemalis* | 0.6084 | | 2.12 | | Not significant | NA |
| *Zonotrichia albicollis* | 0.0856 | | 12.41 | | Not significant | NA |
| *Ficedula albicollis* | 0.3912 | | 0.76 | | Not significant | NA |
| *Acridotheres javanicus* | 0.0197 | | 50.0 | | Not significant | NA |
| *Sturnus vulgaris* | 0.5552 | | 0.68 | | Not significant | NA |
| *Hirundo rustica rustica* | 0.011 | | 8.41 | | Not significant | 0.02725 |
| *Acanthisitta chloris* | 0.137 | | 1.45 | | Not significant | NA |
| *Zosterops lateralis melanops* | 0.3626 | | 0.72 | | Not significant | NA |
| *Lepidothrix coronata* | 0.63 | | 1.55 | | Not significant | NA |
| *Manacus vitellinus* | 0.1007 | | 14.41 | | Not significant | NA |
| *Corvus cornix cornix* | 0.57 | | 2.03 | | Not significant | NA |
| *Corvus brachyrhynchos* | 0.8486 | | 1.12 | | Not significant | NA |
| *Parotia lawesii* | 0.0199 | | 50.00 | | Not significant | NA |
| *Paradisaea raggiana* | 0.0080 | | 0.12/4.9 | | Highly unreliable K | 0.00445 |
| *Pseudopodoces humilis* | 0.6052 | | 0.65 | | Not significant | NA |
| *Parus major* | 0.2419 | | 0.54 | | Not significant | NA |
| *Cyanistes caeruleus* | 0.3529 | | 5.95 | | Not significant | NA |
| *Erythrura gouldiae* | 0.8135 | | 1.02 | | Not significant | NA |
| ***Telluraves*** *(without passerines)* | | | | | | |
| *Leptosomus discolor* | 0.1316 | | 0.58 | | Not significant | NA |
| *Tyto alba* | 0.4337 | | 0.64 | | Not significant | NA |
| *Athene cunicularia* | 0.1581 | | 0.66 | | Not significant | NA |
| *Cariama cristata* | 1.0 | | 1.02/1.0 | | Highly unreliable K | NA |
| *Buceros rhinoceros silvestris* | 0.0441 | | 0.55 | | Not significant | NA |
| *Apaloderma vittatum* | 0.2101 | | 1.45 | | Not significant | NA |
| *Colius striatus* | 0.0092 | | 1.95 | | Not significant | 0.04581 |
| *Aquila chrysaetos canadensis* | 1.0 | | 0.88/1.0 | | Highly unreliable K | NA |
| *Haliaeetus albicilla* | 0.2494 | | 0.0/4.56 | | Highly unreliable K | NA |
| *Haliaeetus leucocephalus* | 1.0 | | 1.19/0.92 | | Highly unreliable K | NA |
| *Falco peregrinus* | 0.9211 | | 1.14 | | Not significant | NA |
| *Falco cherrug* | 0.8097 | | 1.16 | | Not significant | NA |
| *Melopsittacus undulatus* | 0.0732 | | 1.71 | | Not significant | NA |
| *Nestor notabilis* | 0.9389 | | 1.07 | | Not significant | NA |
| ***Aequorlitornithes*** | | | | | | |
| *Uria lomvia* | 0.8404 | | 0.87 | | Not significant | NA |
| *Calidris pygmaea* | 0.1661 | | 2.36 | | Not significant | NA |
| *Calidris pugnax* | 0.3786 | | 0.33 | | Not significant | NA |
| *Pelecanus crispus* | 0.2965 | | 0.07 | | Not significant | NA |
| *Egretta garzetta* | 0.1467 | | 0.43 | | Not significant | NA |
| *Fulmarus glacialis* | 0.5327 | | 1.3 | | Not significant | NA |
| *Nipponia nippon* | 0.026 | | 0.05 | | **Relaxed** | NA |
| *Aptenodytes forsteri* | 0.27 | | 0.18 | | Not significant | NA |
| *Pygoscelis adeliae* | 0.6487 | | 0.47 | | Not significant | NA |
| *Pygoscelis antarcticus* | 0.1619 | | 0.0 | | Not significant | NA |
| *Gavia stellata* | 0.6208 | | 1.29 | | Not significant | NA |
| *Eurypyga helias* | 0.741 | | 0.88 | | Not significant | NA |
| *Phaethon lepturus* | 0.42 | | 2.12 | | Not significant | NA |
| *Phalacrocorax carbo* | 0.8934 | | 1.08 | | Not significant | NA |
| *Urile pelagicus* | 1.0 | | 1.15 | | Not significant | NA |
| *Charadrius vociferus* | 0.0362 | | 16.99 | | **Intensified** | NA |
| ***Galliformes*** | | | | | | |
| *Meleagris gallopavo* | 0.1703 | | 0.0 | | Not significant | NA |
| *Syrmaticus mikado* | 0.347 | | 0.0 | | Not significant | NA |
| *Numida meleagris* | 1.0 | | 0.9/1.9 | | Highly unreliable K | NA |
| *Coturnix japonica* | 0.2803 | | 1.47 | | Not significant | NA |
| *Gallus gallus* | 0.0346 | | 0.0 | | Not significant | NA |
| *Tympanuchus cupido pinnatus* | 0.0 | | 4.95/0.0 | | Highly unreliable K | 0.00184 |
| ***Galliformes* (without *Tympanuchus cupido pinnatus*)** | | | | | | |
| *Meleagris gallopavo* | 0.5280 | | 0.38 | | Not significant | NA |
| *Syrmaticus mikado* | 0.8485 | | 0.07 | | Not significant | NA |
| *Numida meleagris* | 0.1306 | | 0.00/3.04 | | Highly unreliable K | NA |
| *Coturnix japonica* | 0.1144 | | 2.31 | | Not significant | NA |
| *Gallus gallus* | 0.2666 | | 0.00 | | Not significant | NA |
| **Ducks (with *Passeriformes &* chicken *as background species*)** | | | | | | |
| *Anser cygnoides domesticus* | 0.2644 | | 0.79 | | Not significant | NA |
| *Anas platyrhynchos* | 0.016 | | 0.5 | | Not significant | NA |
| *Anas zonorhyncha* | 0.0805 | | 0.62 | | Not significant | NA |
| *Anser brachyrhynchus* | 0.2971 | | 0.81 | | Not significant | NA |
| ***Columbaves*, *Strisores* & *Gruiformes*** | | | | | | |
| *Tauraco erythrolophus* | 0.0383 | | 4.24 | | Not significant | 0.03149 |
| *Balearica regulorum gibbericeps* | 0.3245 | | 0.63 | | Not significant | NA |
| *Cuculus canorus* | 0.0078 | | 34.05 | | Not significant | NA |
| *Caprimulgus carolinensis* | 0.5846 | | 1.15 | | Not significant | NA |
| *Pterocles gutturalis* | 0.4659 | | 1.18 | | Not significant | NA |
| *Chlamydotis macqueenii* | 0.5705 | | 0.83 | | Not significant | NA |
| *Chaetura pelagica* | 0.0017 | | 0.21 | | Not significant | NA |
| *Columba livia* | 0.2325 | | 2.74 | | Not significant | NA |
| *Mesitornis unicolor* | 0.3555 | | 1.19 | | Not significant | NA |

**Supplementary Table S5:** The list of sites identified to be under selection in the cetacean lineage by the programs MEME, FEL, FUBAR and BUSTED from the HyPhy package. Set of species used as the background and test set in the alignment are listed for each of the alignments used.

| **Background species** | **Test species** | **Methods** | **Sites identified as positively selected** |
| --- | --- | --- | --- |
| (Species used in (Endo et al. 2018)) | | | |
| 1. *Camelus bactrianus* 2. *Camelus dromedaries* 3. *Vicugna pacos* 4. *Bison bison bison* 5. *Bos mutus* 6. *Sus scrofa* 7. *Ovis aries* 8. *Capra hircus* | 1. *Balaenoptera acutorostrata scammoni* 2. *Physeter catodon* 3. *Lipotes vexillifer* 4. *Orcinus orca* 5. *Tursiops truncates* | MEME, FEL & FUBAR | (D/N)407(G/S), (N/S)423(D/N) |
|  |  | MEME & FEL | I60(T/A), R73Q, K150(N/S), V161(L/M) |
|  |  | MEME | 554(C-terminal tail) |
|  |  | FEL | F306Y |
|  |  | BUSTED | No Evidence |
| All cetaceans as test species | | | |
| 1. *Capreolus capreolus* 2. *Cervus elaphus hippelaphus* 3. *Elaphurus davidianus* 4. *Odocoileus virginianus texanus* 5. *Bison bison bison* 6. *Bos mutus* 7. *Bos indicus* 8. *Bos taurus* 9. *Bubalus bubalis* 10. *Pantholops hodgsonii* 11. *Capra hircus* 12. *Capra sibirica* 13. *Ovis aries* 14. *Ammotragus lervia* 15. *Giraffa camelopardalis tippelskirchi* 16. *Hippopotamus amphibius* 17. *Sus scrofa* 18. *Vicugna pacos* 19. *Camelus bactrianus* 20. *Camelus dromedarius* | 1. *Balaena mysticetus* 2. *Eschrichtius robustus* 3. *Balaenoptera bonaerensis* 4. *Balaenoptera acutorostrata scammoni* 5. *Balaenoptera physalus* 6. *Lipotes vexillifer* 7. *Lagenorhynchus obliquidens* 8. *Sousa sahulensis* 9. *Sousa chinensis* 10. *Tursiops aduncus* 11. *Tursiops truncatus* 12. *Orcinus orca* 13. *Delphinapterus leucas* 14. *Phocoena phocoena* 15. *Neophocaena asiaeorientalis asiaeorientalis* 16. *Physeter catodon* | MEME,  FEL &  FUBAR | V116(L/M/F/I)  D362(S/N/G) |
|  |  | MEME &  FEL | Q71(L/P)  K105(N/S)  R165H  F261(Y/V)  A294V  S378(N/K/H/D) |
|  |  | MEME | L3V |
|  |  | FEL | I15(T/A) |
| All cetaceans with longer C-terminal tail used as test species | | | |
| 1. *Capreolus capreolus* 2. *Cervus elaphus hippelaphus* 3. *Elaphurus davidianus* 4. *Odocoileus virginianus texanus* 5. *Bison bison bison* 6. *Bos mutus* 7. *Bos indicus* 8. *Bos taurus* 9. *Bubalus bubalis* 10. *Pantholops hodgsonii* 11. *Capra hircus* 12. *Capra sibirica* 13. *Ovis aries* 14. *Ammotragus lervia* 15. *Giraffa camelopardalis tippelskirchi* 16. *Hippopotamus amphibius* 17. *Sus scrofa* 18. *Vicugna pacos* 19. *Camelus bactrianus* 20. *Camelus dromedarius* | 1. *Balaena mysticetus* 2. *Eschrichtius robustus* 3. *Balaenoptera bonaerensis* 4. *Balaenoptera acutorostrata scammoni* 5. *Balaenoptera physalus* 6. *Lipotes vexillifer* 7. *Lagenorhynchus obliquidens* 8. *Orcinus orca* 9. *Delphinapterus leucas* 10. *Phocoena phocoena* 11. *Neophocaena asiaeorientalis asiaeorientalis* 12. *Physeter catodon* | MEME, FEL & FUBAR | V116(L/M/F/I)  D362(S/N/G) |
|  |  | MEME &  FEL | Q71(L/P)  K105(N/S)  F261(Y/V)  S378(N/K/H/D) |
|  |  | MEME | L3V  V7A |
|  |  | FEL | V2M  I15(T/A) |
| All cetaceans with shorter C-terminal tail used as test species | | | |
| 1. *Capreolus capreolus* 2. *Cervus elaphus hippelaphus* 3. *Elaphurus davidianus* 4. *Odocoileus virginianus texanus* 5. *Bison bison bison* 6. *Bos mutus* 7. *Bos indicus* 8. *Bos taurus* 9. *Bubalus bubalis* 10. *Pantholops hodgsonii* 11. *Capra hircus* 12. *Capra sibirica* 13. *Ovis aries* 14. *Ammotragus lervia* 15. *Giraffa camelopardalis tippelskirchi* 16. *Hippopotamus amphibius* 17. *Sus scrofa* 18. *Vicugna pacos* 19. *Camelus bactrianus* 20. *Camelus dromedarius* | 1. *Sousa sahulensis* 2. *Sousa chinensis* 3. *Tursiops aduncus* 4. *Tursiops truncatus* | MEME, FEL & FUBAR | V116(L/M/F/I) |
|  |  | MEME &  FEL | 165  251  294  332 |
|  |  | FUBAR | D362(S/N/G) |

**Supplementary Table S6**: Count of the number of reads in woolly mammoth, American mastodon and straight tusked elephant whole genome sequencing data supporting the bases A, T, G and C at the reading frame disrupting sites in the *CYP8B1* gene on chromosome 19 in the loxAfr4 genome assembly. The number of reads supporting each base was obtained from the bam files using the bam-readcounts utility.

| **Position** | **Reads supporting base** | | | | | **Consensus base** | **Biosample** | **SRA Run Accession** | **Description** |
| --- | --- | --- | --- | --- | --- | --- | --- | --- | --- |
|  | **A** | **T** | **C** | **G** | **Total** |  |  |  |  |
| 27488467 | 0 | **17** | 0 | 0 | 17 | T | SAMEA3340290 | ERR852028 | (*Mammuthus primigenius*)  Oimyakon district, NE Siberia |
| 27488602 | 0 | **10** | 0 | 0 | 10 | T |  |  |  |
| 27489061 | 0 | **8** | 0 | 0 | 8 | T |  |  |  |
| 27489236 | 0 | 0 | **12** | 0 | 12 | C |  |  |  |
| 27489342 | 0 | **7** | 0 | 0 | 7 | T |  |  |  |
| 27488467 | 0 | **17** | 0 | 0 | 17 | T | SAMEA3340289 | ERR855944 | (*Mammuthus primigenius*)  Wrangel Island  (P964) |
| 27488602 | 0 | **13** | 0 | 0 | 13 | T |  |  |  |
| 27489061 | 0 | **16** | 0 | 0 | 16 | T |  |  |  |
| 27489236 | 0 | 0 | **17** | 0 | 17 | C |  |  |  |
| 27489342 | 0 | **15** | 0 | 0 | 15 | T |  |  |  |
| 27488467 | 0 | **3** | 0 | 0 | 3 | T | SAMEA104469184 | ERR2260503 | (*Mammut americanum*)  American mastodon genome from Alaska, USA |
| 27488602 | 0 | **3** | 0 | 0 | 3 | T |  |  |  |
| 27489061 | 0 | **11** | 0 | 0 | 11 | T |  |  |  |
| 27489236 | 0 | 0 | **3** | 0 | 3 | C |  |  |  |
| 27489342 | 0 | **6** | 0 | 0 | 6 | T |  |  |  |
| 27488602 | 0 | **2** | 0 | 0 | 2 | T | SAMEA104469193 | ERR2260508 | (*Mammut americanum*)  American mastodon from Gulf of Maine, USA |
| 27488467 | 0 | **1** | 0 | 0 | 1 | T | SAMEA104469190 | ERR2260505 | (*Mammuthus primigenius*)  Yamal peninsula, Russia |
| 27488602 | 0 | **1** | 0 | 0 | 1 | T |  |  |  |
| 27489236 | 0 | 0 | **2** | 0 | 2 | C |  |  |  |
| 27488602 | 0 | **2** | 0 | 0 | 2 | T | SAMEA104469191 | ERR2260506 | (*Mammuthus columbi*)  Wyoming, USA |
| 27489061 | 0 | **1** | 0 | 0 | 1 | T |  |  |  |
| 27488467 | 0 | **5** | 0 | 0 | 5 | T | SAMEA104469187 | ERR2260504 | (*Elephas antiquus*) Straight-tusked elephant genome from Germany |
| 27488602 | 0 | **5** | 0 | 0 | 5 | T |  |  |  |
| 27489061 | 0 | 0 | 0 | 2 | 2 | G |  |  |  |
| 27489236 | 0 | 0 | **8** | 0 | 8 | C |  |  |  |

**Supplementary Table S7**: Results of performing tests for relaxed selection within the *Chiroptera* order in each of the species included in our multiple sequence alignment after removing one of the species. The K-value represents the intensity of selection. Significant result in the hypothesis test with values of K < 1 refer to relaxed purifying selection and K>1 refers to intensification of selection. None of the significant results survive multiple testing corrections.

| **Test Species** | **p-value** | **K** | **Comment** | **Removed species** |
| --- | --- | --- | --- | --- |
| *Pteropus vampyrus* | 0.0022 | 0 | **Relaxed** | *Desmodus rotundus* |
| *Rhinolophus sinicus* | 0.4196 | 0.24 | Not significant |  |
| *Hipposideros armiger* | 0.0017 | 0/10.28 | Highly unreliable K |  |
| *Eptesicus fuscus* | 0.2099 | 0 | Not significant |  |
| *Myotis davidii* | 0.0852 | 0 | Not significant |  |
| *Myotis lucifugus* | 0.1146 | 10.61 | Not significant |  |
| *Myotis brandtii* | 0.2824 | 0.01 | Not significant |  |
| *Miniopterus natalensis* | 0.0029 | 7.85 | **Intensified** |  |
| *Pteropus vampyrus* | 0.0029 | 0.01 | **Relaxed** | *Eptesicus fuscus* |
| *Pteropus alecto* | 0.0205 | 3.84 | **Intensified** |  |
| *Rhinolophus sinicus* | 0.346 | 0.2 | Not significant |  |
| *Hipposideros armiger* | 0.0023 | 0/10.28 | Highly unreliable K |  |
| *Miniopterus natalensis* | 0.0463 | 1.78 | **Intensified** |  |
| *Myotis davidii* | 0.0527 | 0 | Not significant |  |
| *Myotis lucifugus* | 0.0849 | 0/10.28 | Highly unreliable K |  |
| *Desmodus rotundus* | 0.1675 | 0.07 | Not significant |  |
| *Pteropus vampyrus* | 0.0006 | 0 | **Relaxed** | *Hipposideros armiger* |
| *Pteropus alecto* | 0.0057 | 0/10.28 | Highly unreliable K |  |
| *Rhinolophus sinicus* | 0.7361 | 0.3 | Not significant |  |
| *Desmodus rotundus* | 0.2661 | 0.1 | Not significant |  |
| *Eptesicus fuscus* | 0.2123 | 1.27/10.28 | Highly unreliable K |  |
| *Myotis davidii* | 0.2086 | 1.31/10.28 | Highly unreliable K |  |
| *Myotis lucifugus* | 0.0697 | 0/10.28 | Highly unreliable K |  |
| *Myotis brandtii* | 0.4224 | 1.16/10.28 | Highly unreliable K |  |
| *Miniopterus natalensis* | 0.0281 | 2.15 | **Intensified** |  |
| *Pteropus vampyrus* | 0.0028 | 0.01 | **Relaxed** | *Miniopterus natalensis* |
| *Rhinolophus sinicus* | 0.7923 | 1.25 | Not significant |  |
| *Hipposideros armiger* | 0.0139 | 0.01 | **Relaxed** |  |
| *Myotis davidii* | 0.8595 | 1.26 | Not significant |  |
| *Myotis lucifugus* | 0.1627 | 0/10.28 | Highly unreliable K |  |
| *Desmodus rotundus* | 0.9456 | 1/10.28 | Highly unreliable K |  |
| *Pteropus vampyrus* | 0.0021 | 0.01 | **Relaxed** | *Myotis brandtii* |
| *Pteropus alecto* | 0.017 | 0/10.28 | Highly unreliable K |  |
| *Rhinolophus sinicus* | 0.3797 | 0.23 | Not significant |  |
| *Hipposideros armiger* | 0.001 | 0/10.28 | Highly unreliable K |  |
| *Miniopterus natalensis* | 0.0025 | 7.66 | **Intensified** |  |
| *Eptesicus fuscus* | 0.1133 | 0 | Not significant |  |
| *Myotis lucifugus* | 0.5776 | 0.02 | Not significant |  |
| *Myotis davidii* | 0.113 | 0.01 | Not significant |  |
| *Desmodus rotundus* | 0.1643 | 0.07 | Not significant |  |
| *Pteropus vampyrus* | 0.0026 | 0 | **Relaxed** | *Myotis davidii* |
| *Rhinolophus sinicus* | 0.3294 | 0.25 | Not significant |  |
| *Hipposideros armiger* | 0.0014 | 0.01/10.28 | Highly unreliable K |  |
| *Miniopterus natalensis* | 0.0019 | 8.3 | **Intensified** |  |
| *Eptesicus fuscus* | 0.0766 | 0 | Not significant |  |
| *Myotis lucifugus* | 0.2129 | 13.04 | Not significant |  |
| *Myotis brandtii* | 0.3372 | 0.01 | Not significant |  |
| *Desmodus rotundus* | 0.1364 | 0.06 | Not significant |  |
| *Pteropus vampyrus* | 0.0018 | 0 | **Relaxed** | *Myotis lucifugus* |
| *Pteropus alecto* | 0.0152 | 4.03 | **Intensified** |  |
| *Rhinolophus sinicus* | 0.4329 | 0.27 | Not significant |  |
| *Miniopterus natalensis* | 0.0024 | 7.37 | **Intensified** |  |
| *Myotis brandtii* | 0.1506 | 0 | Not significant |  |
| *Myotis davidii* | 0.0966 | 0 | Not significant |  |
| *Desmodus rotundus* | 0.1609 | 0.06 | Not significant |  |
| *Pteropus vampyrus* | 0.0437 | 0.3 | **Relaxed** | *Pteropus alecto* |
| *Rhinolophus sinicus* | 0.5131 | 1.44/10.28 | Highly unreliable K |  |
| *Hipposideros armiger* | 0.0019 | 0 | **Relaxed** |  |
| *Miniopterus natalensis* | 0.0014 | 10.52 | **Intensified** |  |
| *Eptesicus fuscus* | 0.3142 | 1.18/10.28 | Highly unreliable K |  |
| *Myotis lucifugus* | 0.0859 | 0/10.28 | Not significant |  |
| *Myotis brandtii* | 0.4591 | 0 | Not significant |  |
| *Desmodus rotundus* | 0.3285 | 0.05 | Not significant |  |
| *Pteropus alecto* | 0.1569 | 0.29/10.28 | Highly unreliable K | *Pteropus vampyrus* |
| *Rhinolophus sinicus* | 0.5091 | 0.11 | Not significant |  |
| *Hipposideros armiger* | 0.0003 | 0/10.28 | Highly unreliable K |  |
| *Miniopterus natalensis* | 0.0029 | 10.87 | **Intensified** |  |
| *Myotis lucifugus* | 0.0892 | 0/10.28 | Highly unreliable K |  |
| *Myotis brandtii* | 0.3566 | 0 | Not significant |  |
| *Eptesicus fuscus* | 0.2312 | 0 | Not significant |  |
| *Desmodus rotundus* | 0.2377 | 0.07 | Not significant |  |
| *Pteropus vampyrus* | 0.0022 | 0 | **Relaxed** | *Rhinolophus sinicus* |
| *Pteropus alecto* | 0.0189 | 0/10.28 | Highly unreliable K |  |
| *Hipposideros armiger* | 0.0134 | 0/10.28 | Highly unreliable K |  |
| *Desmodus rotundus* | 0.1863 | 0.09 | Not significant |  |
| *Eptesicus fuscus* | 0.1272 | 0 | Not significant |  |
| *Myotis davidii* | 0.0764 | 0.01 | Not significant |  |
| *Myotis lucifugus* | 0.0842 | 0/10.28 | Highly unreliable K |  |
| *Myotis brandtii* | 0.2484 | 0 | Not significant |  |
| *Miniopterus natalensis* | 0.0034 | 10.03 | **Intensified** |  |
| *Rhinolophus sinicus* | 1 | 1.04/10.28 | Highly unreliable K | *Rousettus aegyptiacus* |
| *Hipposideros armiger* | 0.0382 | 0 | **Relaxed** |  |
| *Miniopterus natalensis* | 0.0297 | 1.85 | **Intensified** |  |
| *Eptesicus fuscus* | 0.7659 | 1.29 | Not significant |  |
| *Myotis davidii* | 0.4935 | 1.34 | Not significant |  |
| *Myotis brandtii* | 1 | 1.25/10.28 | Highly unreliable K |  |
| *Myotis lucifugus* | 0.1222 | 0 | Not significant |  |
| *Desmodus rotundus* | 1 | 0.97/10.28 | Highly unreliable K |  |
| *Pteropus vampyrus* | 0.0026 | 0 | **Relaxed** |  |

**Supplementary Table S8**: Raw read support for correction of the *CYP8B1* gene assembly in *Myotis lucifugus*, *Serinus canaria*.

| **SRA Experiments** | **Isolate** | **Species** | **Reads supporting correction** |
| --- | --- | --- | --- |
| SRX2991344, SRX2991343,  SRX2991351,  SRX2991352 | PAGC60639 | *Myotis lucifugus* | SRA:SRR5812836.28289254.1  SRA:SRR5812837.36405657.1  SRA:SRR5812837.34855292.1  SRA:SRR5812837.30302084.1  SRA:SRR5812837.10537349.1  SRA:SRR5812837.1091412.1  SRA:SRR5812836.11141664.1  SRA:SRR5812828.33758701.1 |
| SRX2991345,  SRX2991346,  SRX2991347,  SRX2991348 | PAGC60619 |  | SRA:SRR5812835.34529646.1  SRA:SRR5812835.29684147.1 |
| SRX2991349,  SRX2991350,  SRX2991361,  SRX2991362 | PAGC60616 |  | SRA:SRR5812830.11161915.1  SRA:SRR5812819.38183257.1 |
| SRX2991357,  SRX2991358,  SRX2991359,  SRX2991360 | PAGC60606 |  | SRA:SRR5812822.29731726.1  SRA:SRR5812821.33715966.1  SRA:SRR5812821.19581651.1  SRA:SRR5812821.17922732.1 |
| SRX2991363,  SRX2991364,  SRX2991365,  SRX2991366 | PAGC60645 |  | SRA:SRR5812815.14797064.1  SRA:SRR5812814.48366008.1 |
| SRX365372 | SAMN02377925 |  | SRA:SRR1013468.4460525.1 |
| SRX326824  SRX326826  SRX326825 | SAMN02256316  SAMN02256322  SAMN02256321 | *Ursus arctos* | SRA: SRR941812.2174683.2  SRA: SRR941811.20468787.1  SRA: SRR941810.18626508.2  SRA: SRR941813.27140493.1 |
| SRX1412949 | SAMN04225736 | *Serinus canaria* | SRA:SRR2896023.111950107.2  SRA:SRR2896023.89584557.1  SRA:SRR2896023.89133431.1  SRA:SRR2896023.29790993.1  SRA:SRR2896023.74795315.2  SRA:SRR2896023.99632397.1  SRA:SRR2896023.28133309.1  SRA:SRR2896023.36108831.1  SRA:SRR2896023.81670045.1  SRA:SRR2896023.29305928.1  SRA:SRR2896023.107190545.1  SRA:SRR2896023.58774899.1  SRA:SRR2896023.53560221.1  SRA:SRR2896023.37753905.1  SRA:SRR2896023.48729682.2 |

**Supplementary Table S9**: List of species whose *CYP8B1* gene was re-annotated leading to a single exon open reading frame of approximately 500 amino acids.

| **Species** | **Transcript ID** | **Exon count** | **ORF length**  (amino acid count) | |
| --- | --- | --- | --- | --- |
|  |  |  | Original | Corrected |
| Tarsier (*Tarsius syrichta*) | ENSTSYT00000036012.1 | 3 | 493 | 501 |
| Sooty mangabey (*Cercocebus atys*) | ENSCATT00000049888.1 | 3 | 495 | 501 |
| Ferret (*Mustela putorius furo*) | ENSMPUT00000019478.1 | 1 | 499 | 501 |
| Cow (*Bos Taurus*) | ENSBTAT00000048363.3 | 2 | 166 | 499 |
| Goat (*Capra hircus*) | ENSCHIT00000038195.1 | 2 | 498 | 499 |
| Marmoset (*Callithrix jacchus*) | ENSCJAT00000003214.3 | 3 | 495 | 501 |
| Bolivian squirrel monkey (*Saimiri boliviensis boliviensis*) | ENSSBOT00000032004.1 | 3 | 495 | 501 |
| Chimpanzee (*Pan troglodytes*) | ENSPTRT00000027673.4 | 2 | 496 | 501 |
| Angola colobus (*Colobus angolensis palliatus*) | ENSCANT00000044551.1 | 2 | 496 | 501 |
| Olive baboon (*Papio anubis*) | ENSPANT00000001183.2 | 2 | 496 | 501 |
| Bonobo (*Pan paniscus*) | ENSPPAT00000051436.1 | 2 | 496 | 501 |
| Black snub-nosed monkey (*Rhinopithecus bieti*) | ENSRBIT00000051574.1 | 2 | 496 | 501 |
| Capuchin (*Cebus capucinus imitator*) | ENSCCAT00000025292.1 | 2 | 496 | 501 |
| Gorilla (*Gorilla gorilla gorilla*) | ENSGGOG00000011191 | 2 | 496 | 501 |
| Golden snub-nosed monkey (*Rhinopithecus roxellana*) | ENSRROT00000030381.1 | 2 | 496 | 501 |
| Crab-eating macaque (*Macaca fascicularis*) | ENSMFAT00000017804.1 | 2 | 496 | 501 |
| Pig-tailed macaque (*Macaca nemestrina*) | ENSMNET00000056089.1 | 2 | 496 | 501 |
| Drill (*Mandrillus leucophaeus*) | ENSMLET00000055398.1 | 2 | 496 | 501 |

**Supplementary Table S10**: Count of the number of reads supporting the bases A, T, G and C at the position *2105805* on chromosome 2 in the Gallus_gallus_5 genome assembly. The number of reads supporting each base was obtained from the bam files using the bam-readcounts utility.

| **SRA Run Accession** | **Reads supporting base** | | | | | **Consensus base** | **Biosample** | **Breed Description** |
| --- | --- | --- | --- | --- | --- | --- | --- | --- |
|  | **A** | **T** | **C** | **G** | **Total** |  |  |  |
| SRR3954707 | 0 | **31** | 0 | 0 | 31 | T | SAMN02981218 | Red Jungle Fowl, inbred line UCD001, RJF #256 (female) |
| SRR1217539 | **14** | **10** | 0 | 0 | 24 | A/T | SAMN02712033 | Yunnan chicken |
| SRR1217491 | **25** | 0 | 0 | 0 | 25 | A | SAMN02712045 | Tibet chicken |
| SRR1217493 | **13** | 0 | 0 | 0 | 13 |  | SAMN02712047 |  |
| SRR1217494 | **16** | 0 | 0 | 1 | 17 |  | SAMN02712048 |  |
| SRR1217496 | **14** | 0 | 0 | 0 | 14 |  | SAMN02712049 |  |
| SRR1217500 | **10** | 0 | 1 | 0 | 11 |  | SAMN02712051 |  |
| SRR1217502 | **43** | 0 | 0 | 0 | 43 |  | SAMN02712052 |  |
| SRR1217507 | **11** | 0 | 0 | 0 | 11 |  | SAMN02712054 |  |
| SRR1217524 | **18** | 0 | 0 | 0 | 18 |  | SAMN02712039 | Red Jungle Fowl |
| SRR1217527 | **35** | 0 | 0 | 0 | 35 |  | SAMN02712040 |  |
| SRR1217529 | **21** | 0 | 0 | 0 | 21 |  | SAMN02712041 |  |
| SRR1217531 | **20** | 0 | 0 | 0 | 20 |  | SAMN02712042 |  |
| SRR1217533 | **35** | 0 | 0 | 0 | 35 |  | SAMN02712043 |  |
| SRR1217535 | **17** | 0 | 1 | 0 | 18 |  | SAMN02712031 | Yunnan chicken |
| SRR1217537 | **36** | 0 | 0 | 0 | 36 |  | SAMN02712032 |  |
| SRR1217542 | **42** | 0 | 0 | 0 | 42 |  | SAMN02712034 |  |
| SRR1217545 | **24** | 0 | 0 | 0 | 24 |  | SAMN02712035 |  |
| SRR1217547 | **13** | 0 | 0 | 0 | 13 |  | SAMN02712036 |  |
| SRR1217549 | **14** | 0 | 0 | 0 | 14 |  | SAMN02712037 |  |
| SRR1217551 | **71** | 0 | 0 | 0 | 71 |  | SAMN02712038 |  |
| SRR2131198 | **19** | 0 | 0 | 0 | 19 |  | SAMN03940091 | White leghorn |
| SRR2131199 | **17** | 0 | 0 | 0 | 17 |  | SAMN03940092 |  |
| SRR2131201 | **16** | 0 | 0 | 0 | 16 |  | SAMN03940093 |  |
| SRR2131205 | **19** | 0 | 0 | 0 | 19 |  | SAMN03940099 | Araucana |
| SRR2131208 | **10** | 0 | 0 | 0 | 10 |  | SAMN03940098 |  |
| SRR3036360 | **13** | 0 | 0 | 0 | 13 |  | SAMN04349707 | Miyi fowl (LCMY6) |
| SRR3041115 | **38** | 0 | 1 | 0 | 39 |  |  | Emei black fowl (LCEM2) |
| SRR3041116 | **15** | 1 | 0 | 0 | 16 |  |  | Emei black fowl (LCEM3) |
| SRR3041121 | **69** | 1 | 0 | 0 | 70 |  |  | Emei black fowl (LCEM4) |
| SRR3041122 | **49** | 0 | 0 | 0 | 49 |  |  | Emei black fowl (LCEM5) |
| SRR3041123 | **15** | 0 | 0 | 0 | 15 |  |  | Emei black fowl (LCEM6) |
| SRR3041124 | **11** | 0 | 0 | 0 | 11 |  |  | Jiuyuan black-bone fowl (LCJY1) |
| SRR3041125 | **63** | 0 | 0 | 0 | 63 |  |  | Jiuyuan black-bone fowl (LCJY3) |
| SRR3041126 | **18** | 0 | 0 | 0 | 18 |  |  | Jiuyuan black-bone fowl (LCJY5) |
| SRR3041127 | **20** | 0 | 0 | 0 | 20 |  |  | Jiuyuan black-bone fowl (LCJY7) |
| SRR3041128 | **20** | 0 | 0 | 0 | 20 |  |  | Jiuyuan black-bone fowl (LCJY8) |
| SRR3041129 | **20** | 0 | 0 | 0 | 20 |  |  | Jinyang silky fowl (LCLS1) |
| SRR3041130 | **22** | 0 | 0 | 0 | 22 |  |  | Jinyang silky fowl (LCLS2) |
| SRR3041131 | **22** | 0 | 0 | 0 | 22 |  |  | Jinyang silky fowl (LCLS3) |
| SRR3041132 | **17** | 0 | 0 | 0 | 17 |  |  | Jinyang silky fowl (LCLS4) |
| SRR3041133 | **17** | 0 | 0 | 0 | 17 |  |  | Jinyang silky fowl (LCLS5) |
| SRR3041135 | **15** | 0 | 0 | 0 | 15 |  |  | Muchuan black-bone fowl (LCMC1) |
| SRR3041136 | **25** | 1 | 0 | 0 | 26 |  |  | Muchuan black-bone fowl (LCMC2) |
| SRR3041137 | **56** | 0 | 0 | 0 | 56 |  |  | Muchuan black-bone fowl (LCMC4) |
| SRR3041138 | **20** | 0 | 0 | 0 | 20 |  |  | Muchuan black-bone fowl (LCMC5) |
| SRR3041364 | **20** | 0 | 0 | 0 | 20 |  |  | Muchuan black-bone fowl (LCMC8) |
| SRR3041409 | **46** | 0 | 0 | 0 | 46 |  |  | Miyi fowl (LCMY1) |
| SRR3041410 | **79** | 1 | 0 | 1 | 81 |  |  | Miyi fowl (LCMY2) |
| SRR3041411 | **60** | 0 | 0 | 0 | 60 |  |  | Miyi fowl (LCMY3) |
| SRR3041412 | **36** | 0 | 0 | 1 | 37 |  |  | Miyi fowl (LCMY4) |
| SRR3041413 | **14** | 0 | 0 | 0 | 14 |  |  | Pengxian yellow fowl (LCPX1) |
| SRR3041414 | **26** | 0 | 0 | 0 | 26 |  |  | Pengxian yellow fowl (LCPX2) |
| SRR3041415 | **16** | 1 | 0 | 0 | 17 |  |  | Pengxian yellow fowl (LCPX3) |
| SRR3041417 | **24** | 0 | 0 | 0 | 24 |  |  | Pengxian yellow fowl (LCPX5) |
| SRR3041418 | **19** | 0 | 0 | 0 | 19 |  |  | Pengxian yellow fowl (LCPX6) |
| SRR3041419 | **23** | 0 | 0 | 0 | 23 |  |  | Shimian caoke fowl (LCSM4) |
| SRR3041420 | **19** | 0 | 0 | 0 | 19 |  |  | Shimian caoke fowl (LCSM5) |
| SRR3041421 | **28** | 0 | 0 | 0 | 28 |  |  | Shimian caoke fowl (LCSM6) |
| SRR3041422 | **19** | 0 | 0 | 0 | 19 |  |  | Shimian caoke fowl (LCSM8) |
| SRR3041423 | **30** | 0 | 0 | 0 | 30 |  |  | Tianfu black-bone fowl (LCTF2) |
| SRR3041425 | **91** | 0 | 0 | 0 | 91 |  |  | Tianfu black-bone fowl (LCTF3) |
| SRR3041426 | **19** | 0 | 0 | 1 | 20 |  |  | Tianfu black-bone fowl (LCTF4) |
| SRR3041427 | **14** | 0 | 0 | 0 | 14 |  |  | Tianfu black-bone fowl (LCTF6) |
| SRR3041428 | **15** | 0 | 0 | 0 | 15 |  |  | Tianfu black-bone fowl (LCTF8) |
| SRR3041433 | **16** | 0 | 0 | 0 | 16 |  |  | Tibetan fowl (TCAB1) |
| SRR3041434 | **31** | 0 | 0 | 0 | 31 |  |  | Tibetan fowl (TCAB3) |
| SRR3041435 | **14** | 0 | 0 | 0 | 14 |  |  | Tibetan fowl (TCAB5) |
| SRR3041436 | **76** | 0 | 0 | 1 | 77 |  |  | Tibetan fowl (TCAB6) |
| SRR3041437 | **18** | 0 | 0 | 0 | 18 |  |  | Tibetan fowl (TCAB7) |
| SRR3041438 | **21** | 0 | 0 | 0 | 21 |  |  | Tibetan fowl (TCDQ1) |
| SRR3041439 | **22** | 0 | 0 | 0 | 22 |  |  | Tibetan fowl (TCDQ2) |
| SRR3041440 | **33** | 0 | 0 | 0 | 33 |  |  | Tibetan fowl (TCDQ3) |
| SRR3041441 | **53** | 0 | 1 | 0 | 54 |  |  | Tibetan fowl (TCDQ4) |
| SRR3041442 | **23** | 0 | 0 | 0 | 23 |  |  | Tibetan fowl (TCDQ5) |
| SRR3041443 | **77** | 0 | 0 | 0 | 77 |  |  | Tibetan fowl (TCDQ6) |
| SRR3041444 | **18** | 0 | 0 | 0 | 18 |  |  | Tibetan fowl (TCGZ1) |
| SRR3041445 | **22** | 0 | 0 | 0 | 22 |  |  | Tibetan fowl (TCGZ10) |
| SRR3041446 | **38** | 0 | 0 | 0 | 38 |  |  | Tibetan fowl (TCGZ3) |
| SRR3041447 | **23** | 0 | 0 | 0 | 23 |  |  | Tibetan fowl (TCGZ4) |
| SRR3041448 | **58** | 0 | 0 | 0 | 58 |  |  | Tibetan fowl (TCGZ5) |
| SRR3041449 | **63** | 0 | 0 | 0 | 63 |  |  | Tibetan fowl (TCGZ6) |
| SRR3041450 | **22** | 0 | 0 | 0 | 22 |  |  | Tibetan fowl (TCLZ1) |
| SRR3041451 | **27** | 0 | 0 | 0 | 27 |  |  | Tibetan fowl (TCLZ2) |
| SRR3041453 | **41** | 0 | 0 | 0 | 41 |  |  | Tibetan fowl (TCLZ4) |
| SRR3041454 | **33** | 1 | 0 | 0 | 34 |  |  | Tibetan fowl (TCLZ5) |
| SRR3041455 | **14** | 0 | 0 | 0 | 14 |  |  | Tibetan fowl (TCQH1) |
| SRR3041456 | **14** | 0 | 0 | 0 | 14 |  |  | Tibetan fowl (TCQH10) |
| SRR3041457 | **20** | 0 | 0 | 0 | 20 |  |  | Tibetan fowl (TCQH11) |
| SRR3041458 | **14** | 0 | 0 | 0 | 14 |  |  | Tibetan fowl (TCQH5) |
| SRR3041504 | **21** | 0 | 0 | 0 | 21 |  |  | Tibetan fowl (TCQH8) |
| SRR3041573 | **24** | 0 | 0 | 0 | 24 |  |  | Tibetan fowl (TCQH9) |
| SRR3041620 | **22** | 0 | 0 | 0 | 22 |  |  | Tibetan fowl (TCSN1) |
| SRR3041692 | **34** | 0 | 0 | 0 | 34 |  |  | Tibetan fowl (TCSN3) |
| SRR3041713 | **38** | 0 | 0 | 0 | 38 |  |  | Tibetan fowl (TCSN4) |
| SRR3041781 | **33** | 0 | 0 | 0 | 33 |  |  | Tibetan fowl (TCSN5) |
| SRR3041923 | **22** | 0 | 0 | 0 | 22 |  |  | Tibetan fowl (TCSN6) |
| SRR3041924 | **11** | 0 | 0 | 0 | 11 |  |  | Tibetan fowl (TCSN7) |
| SRR3041925 | **19** | 0 | 0 | 0 | 19 |  |  | Tibetan fowl (TCSN8) |
| SRR3041926 | **18** | 0 | 0 | 0 | 18 |  |  | Tibetan fowl (TCSN9) |

**Supplementary Figure legends**

**Supplementary Figure S1:** A pixel plot comparing the multiple sequence alignments of the *CYP8B1* gene generated by four methods: CLUSTALW, MAFFT, MUSCLE and PRANK.

**Supplementary Figure S2:** GC deviation and GC content are calculated in windows of size 100 with a step size of 10 for each of the species. **(A)** GC deviation and **(B)** GC content are visualised for each group of species.

**Supplementary Figure S3: (A)** Signatures of relaxed and intensified selection inferred by RELAX general descriptive model across artiodactyl species. **(B)** Signatures of relaxed and intensified selection inferred by RELAX general descriptive model across Afrotheria species. **(C)** Signatures of relaxed and intensified selection inferred by RELAX general descriptive model across carnivore species.

**Supplementary Figure S4:** Signatures of relaxed selection represented as a combination of K values (K<1 is relaxed selection & K>1 intensified selection) denoting the intensity of change in selection and corresponding p-values in *Galliformes* species. Each data point in the figure is the result of one hypothesis test each*.* Tests that showed significant relaxation of selection are shown in filled squares while those that are not significant are shown as empty circles. The colour of each data point corresponds to one of the three tree topologies shown in panel A. **(A)** Tree topologies used to assess signatures of relaxed selection **(B, C)** Each of the six *Galliformes* species tested.
