## Supplementary Figures for "Signatures of relaxed selection in the *CYP8B1* gene of birds and mammals"

### Slide 1
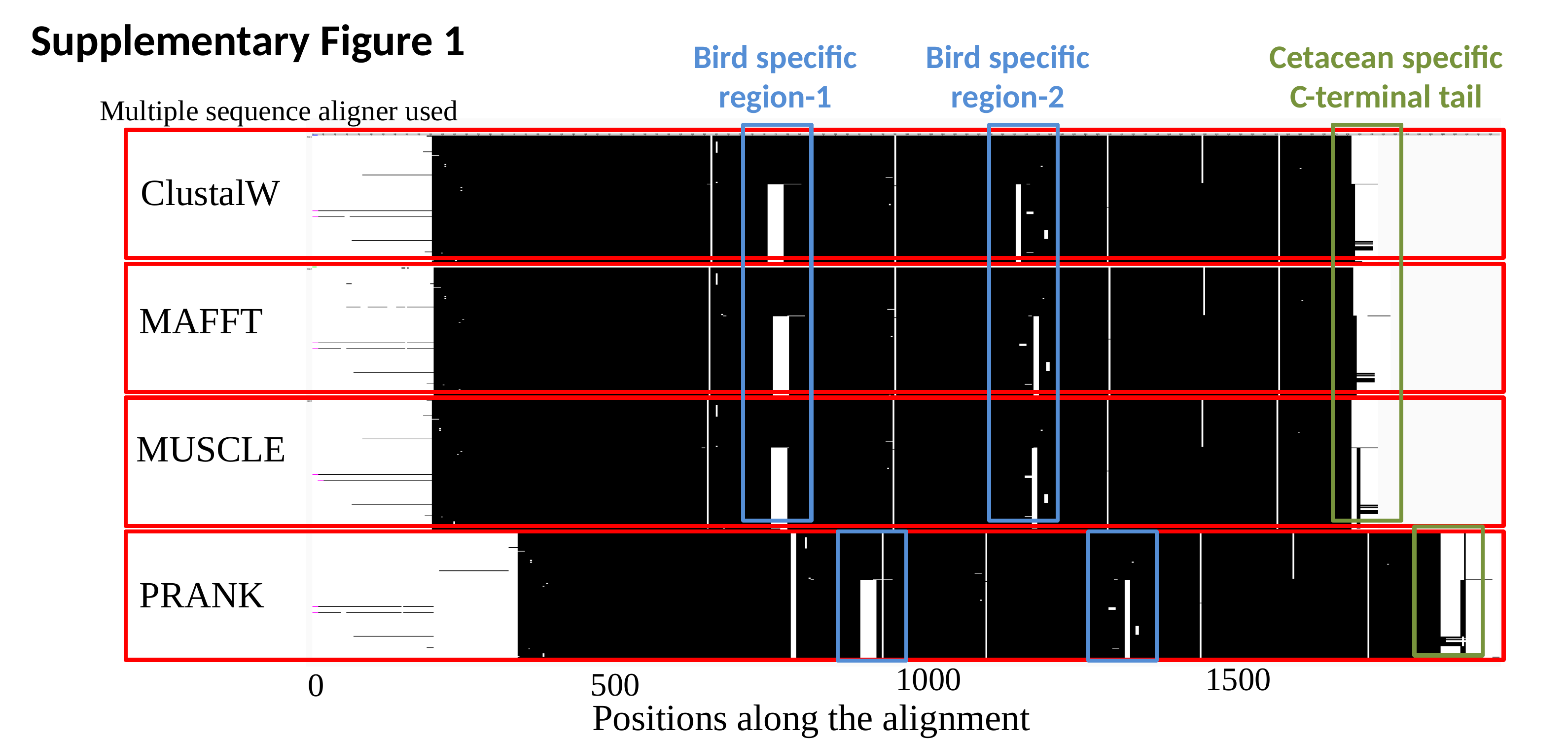

Supplementary Figure 1
Bird specific
region-1
Cetacean specific
C-terminal tail
Bird specific
region-2
Multiple sequence aligner used
ClustalW
MAFFT
MUSCLE
PRANK
1000
1500
500
0
Positions along the alignment

### Slide 2
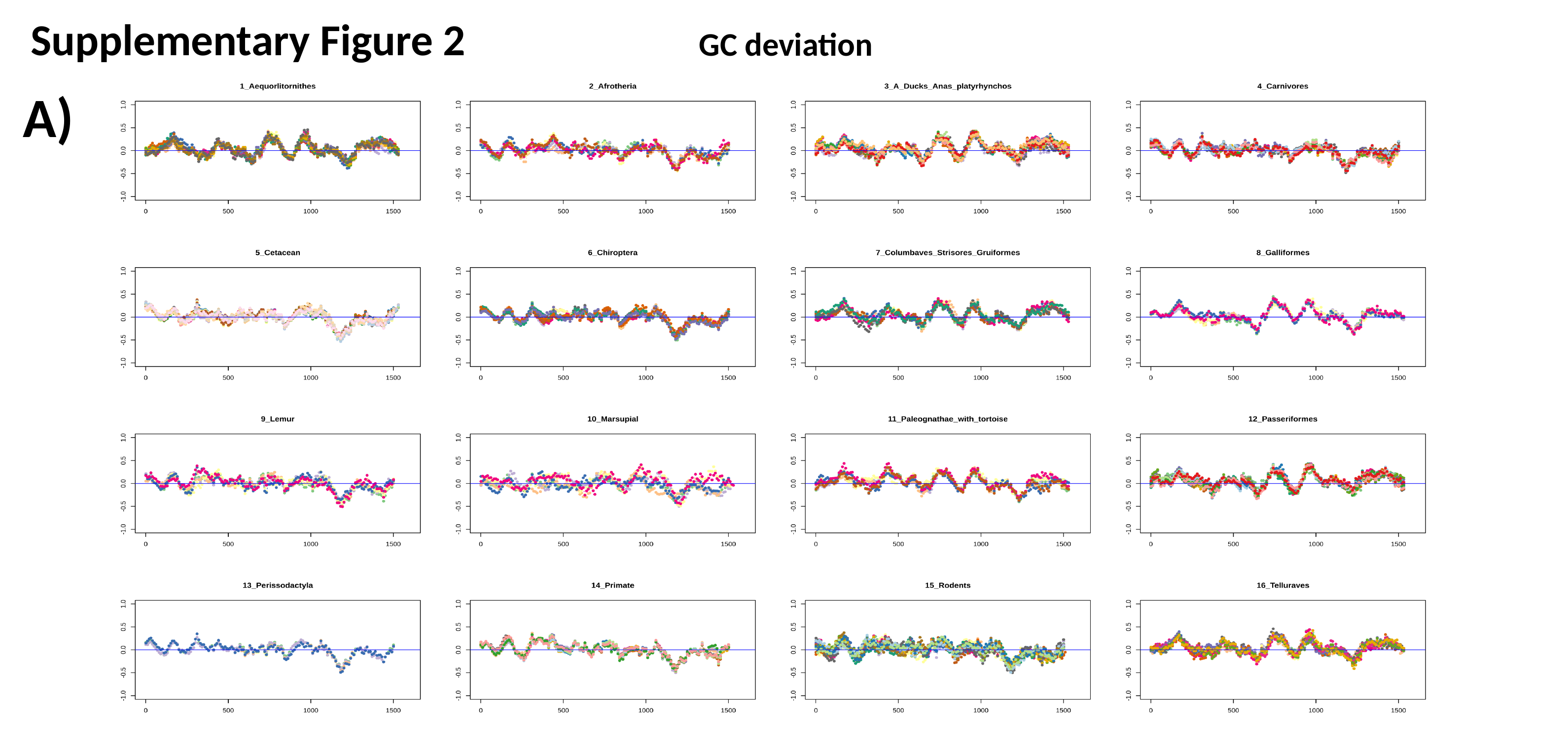

Supplementary Figure 2
GC deviation
A)

### Slide 3
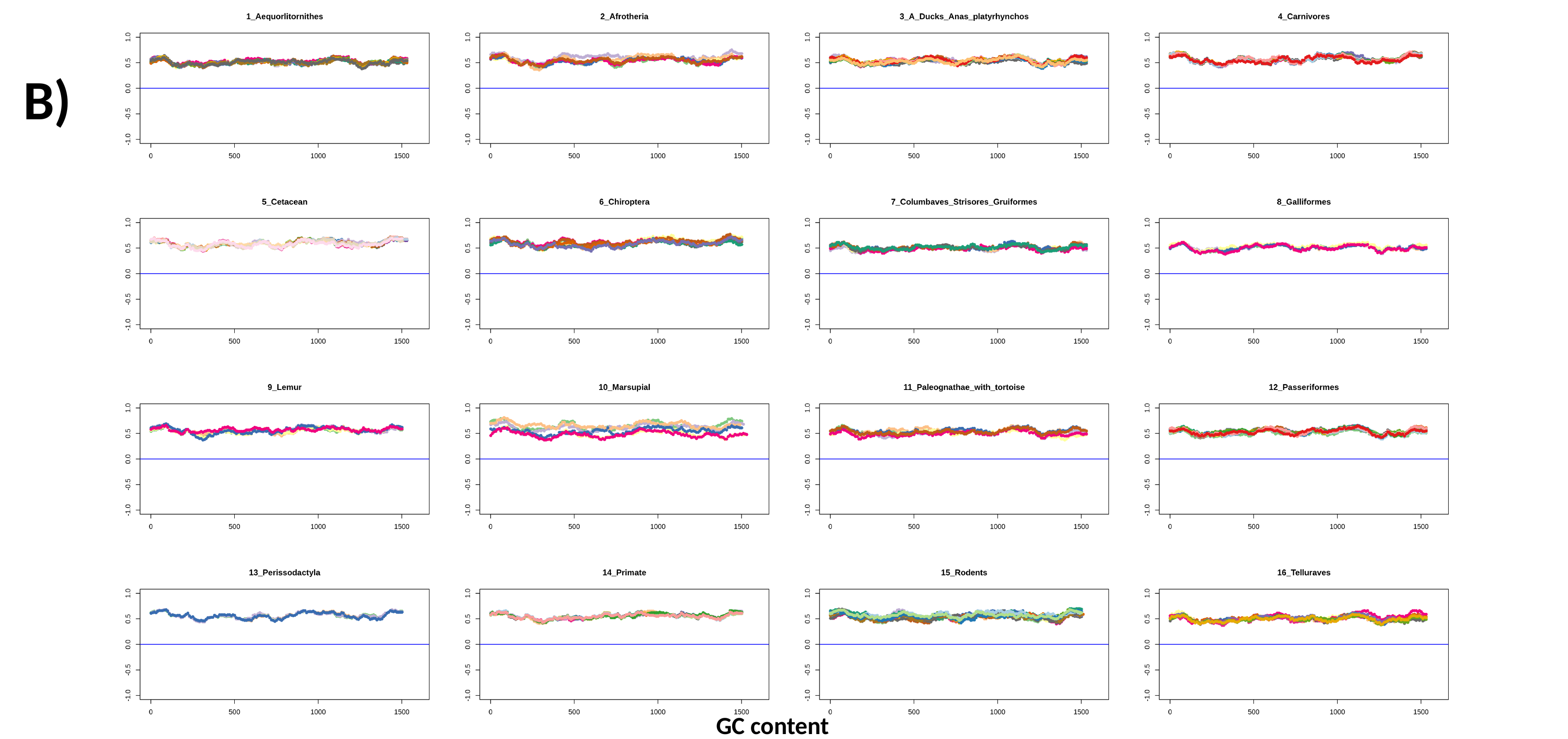

B)
GC content

### Slide 4
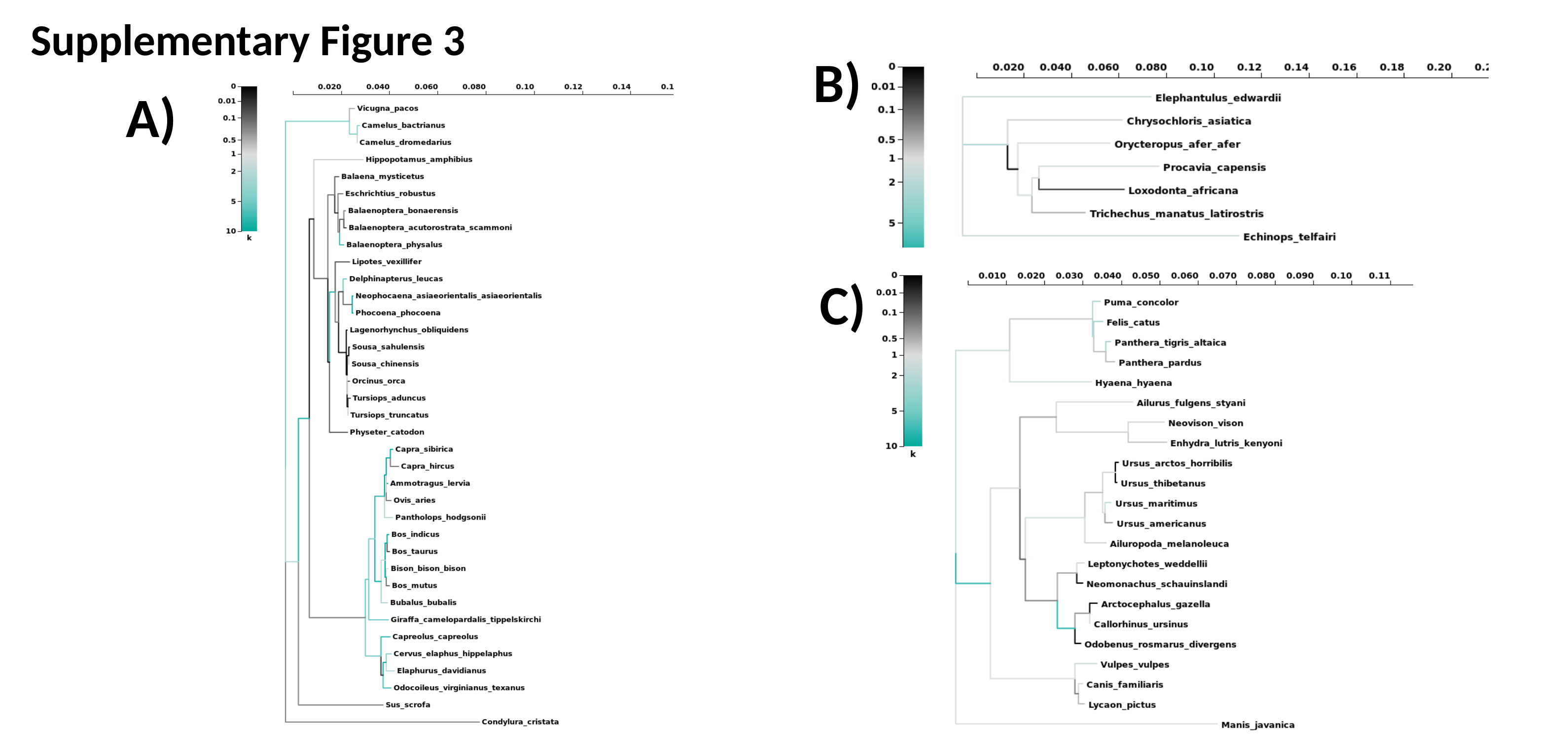

Supplementary Figure 3
B)
A)
C)

### Slide 5
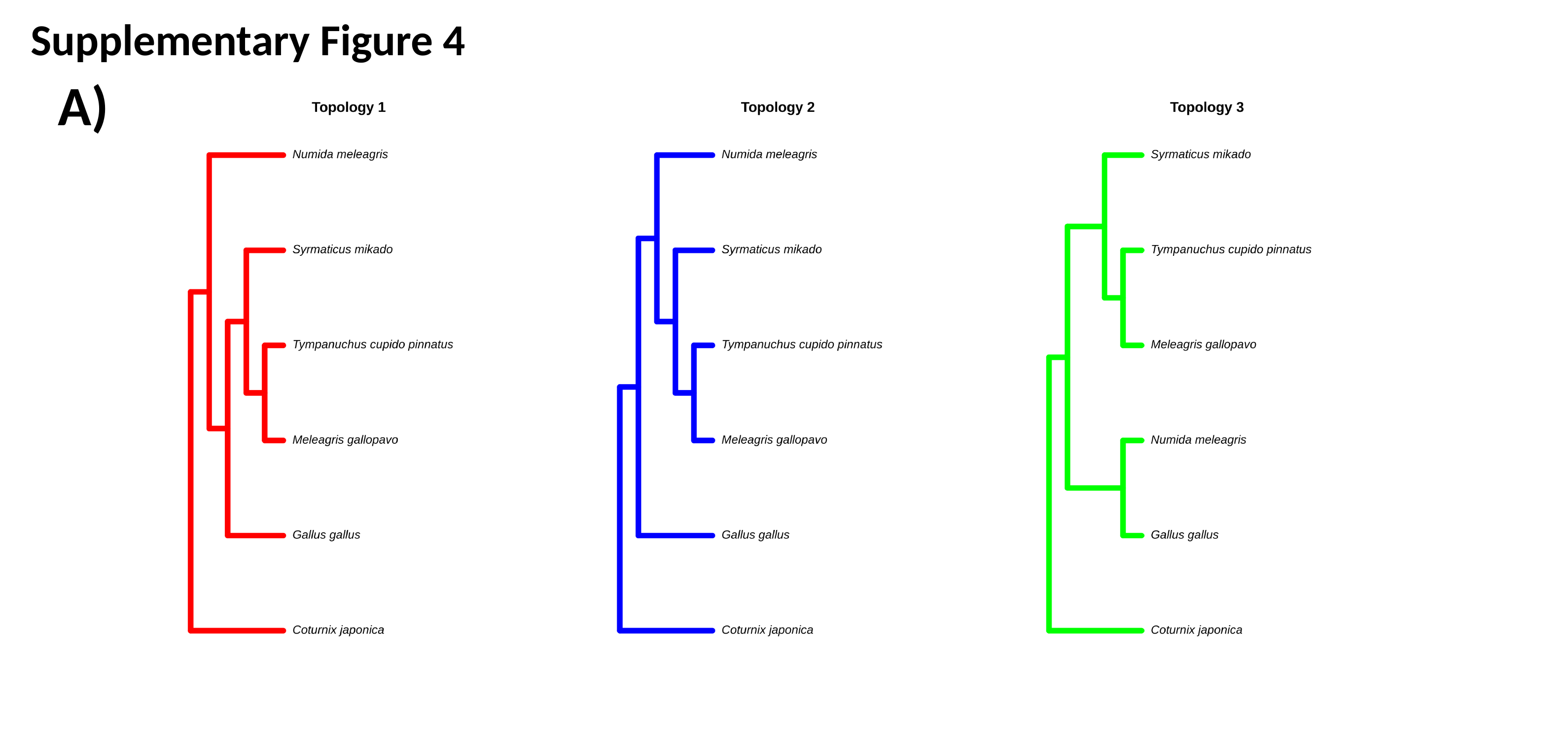

Supplementary Figure 4
A)

### Slide 6
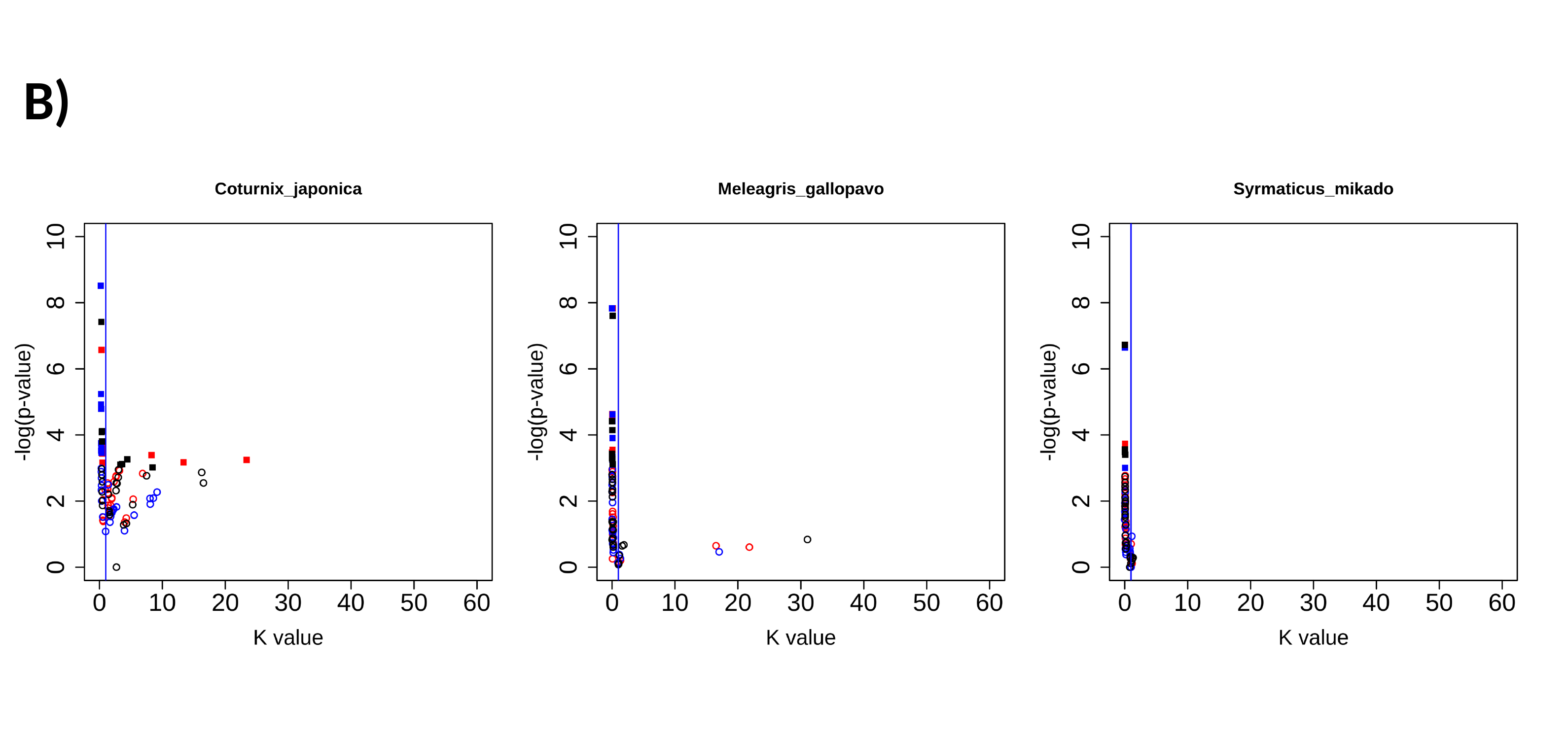

B)

### Slide 7
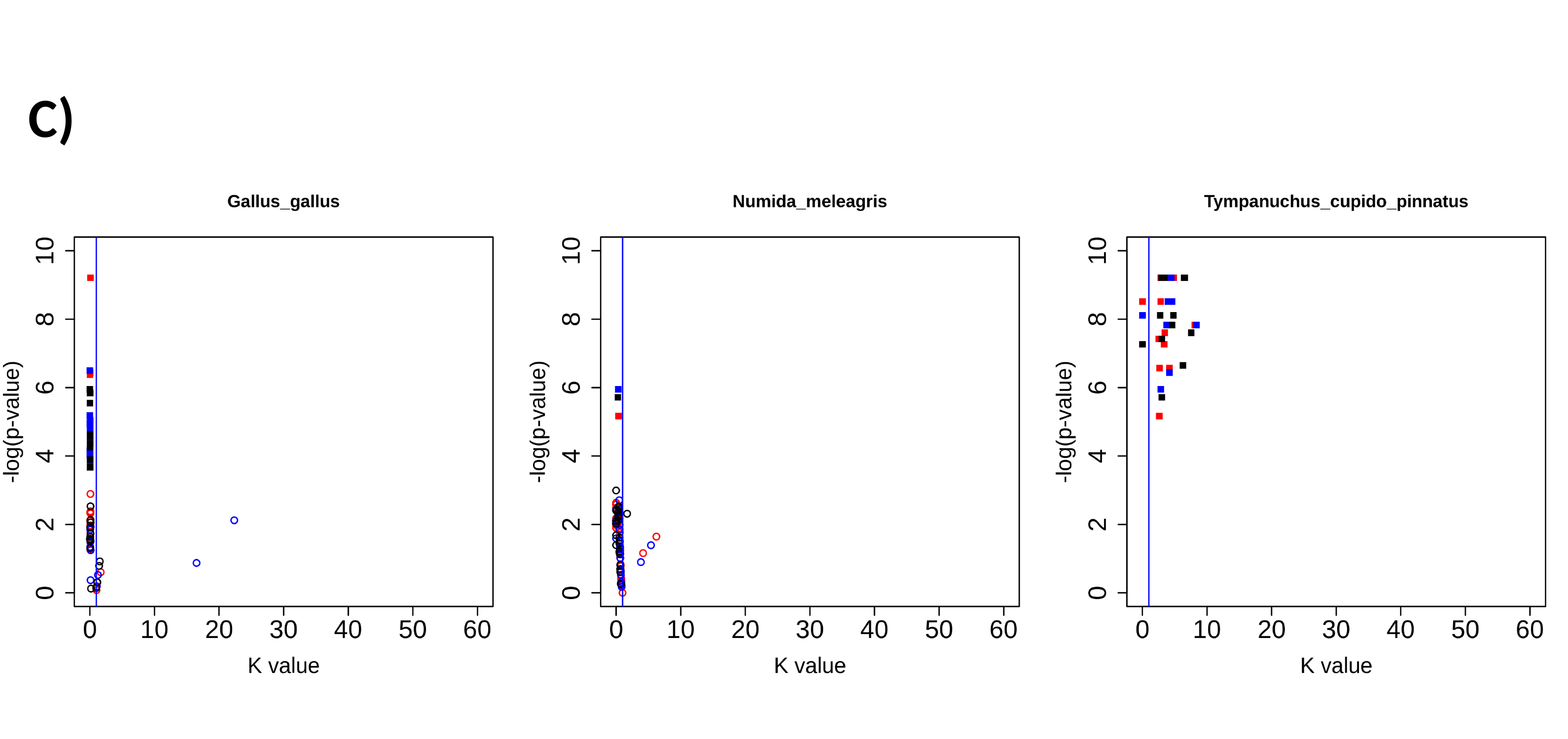

C)
